# Loss of *slc39a14* causes simultaneous manganese deficiency and hypersensitivity in zebrafish

**DOI:** 10.1101/2020.01.31.921130

**Authors:** Karin Tuschl, Richard J White, Leonardo E Valdivia, Stephanie Niklaus, Isaac H Bianco, Ian M Sealy, Stephan CF Neuhauss, Corinne Houart, Stephen W Wilson, Elisabeth M Busch-Nentwich

**Affiliations:** Department of Cell and Developmental Biology, University College London, Gower Street, WC1E 6BT, UK; Department of Developmental Neurobiology and MRC Centre for Neurodevelopmental Disorders, loPPN, Kings College London, New Hunt’s House, Guy’s Campus, London, SE1 1UL, UK; UCL GOS Institute of Child Health, 30 Guilford Street, London, WC1N 1EH, UK; Wellcome Sanger Institute, Wellcome Genome Campus, CB10 1SA, UK; Cambridge Institute of Therapeutic Immunology & Infectious Disease (CITIID), Jeffrey Cheah Biomedical Centre, University of Cambridge, Puddicombe Way, Cambridge, CB2 0AW; Center for Integrative Biology, Facultad de Ciencias, Universidad Mayor, Santiago, Chile; Institute of Molecular Life Sciences, University of Zurich, Winterthurerstrasse 190, 8057, Zurich, Switzerland; Department of Neuroscience, Physiology & Pharmacology, University College London, Gower Street, WC1E 6BT, UK

**Keywords:** *slc39a14*, manganese, transcriptome, neurotoxicity, vision, calcium

## Abstract

Mutations in SLC39A14, a manganese uptake transporter, lead to a neurodegenerative disorder characterised by accumulation of manganese in the brain and rapidly progressive dystonia-parkinsonism (Hypermanganesemia with Dystonia 2, HMNDYT2). Similar to the human phenotype, zebrafish *slc39a14^U801-/-^* mutants show prominent brain manganese accumulation and abnormal locomotor behaviour. In order to identify novel potential targets of manganese neurotoxicity, we performed transcriptome analysis of individual homozygous mutant and sibling *slc39a14^U801^* zebrafish at five days post fertilisation unexposed and exposed to MnCl_2_. Anatomical gene enrichment analysis confirmed that differentially expressed genes map to the central nervous system and eye. Biological interpretation of differentially expressed genes suggests that calcium dyshomeostasis, activation of the unfolded protein response, oxidative stress, mitochondrial dysfunction, lysosomal disruption, apoptosis and autophagy, and interference with proteostasis are key events in manganese neurotoxicity. Differential expression of visual phototransduction genes also predicted visual dysfunction in mutant larvae which was confirmed by the absence of visual background adaptation and a diminished optokinetic reflex. Surprisingly, we found a group of differentially expressed genes in mutant larvae that normalised upon MnCl_2_ treatment suggesting that, in addition to neurotoxicity, manganese deficiency is present either subcellularly or in specific cells or tissues. This may have important implications for treatment as manganese chelation may aggravate neurological symptoms. Our analyses show that *slc39a14^U801-/-^* mutant zebrafish present a powerful model to study the cellular and molecular mechanisms underlying disrupted manganese homeostasis.

**Significance statement:** Manganese neurotoxicity leading to progressive dystonia-parkinsonism is a characteristic feature of Hypermanganesemia with dystonia 2 (HMNDYT2) caused by mutations in SLC39A14, a manganese uptake transporter. Transcriptional profiling in *slc39a14^U801^* loss-of-function zebrafish suggests that, in addition to manganese neurotoxicity, subcellular or cell type specific manganese deficiency contributes to the disease phenotype. Both manganese overload and deficiency appear to be associated with Ca^2+^ dyshomeostasis. We further demonstrate that activation of the unfolded protein response, oxidative stress, mitochondrial dysfunction, apoptosis and autophagy, and disrupted proteostasis are likely downstream events in manganese neurotoxicity. Our study shows that the zebrafish *slc39a14^U801^* loss-of-function mutant is a powerful model to elucidate the mechanistic basis of diseases affected by manganese dyshomeostasis.

## Introduction

SLC39A14 is a manganese (Mn) uptake transporter essential for the maintenance of Mn homeostasis (Thompson and Wessling-Resnick, 2019). Mutations in SLC39A14 impair cellular Mn uptake and result in systemic Mn overload characterised by significant hypermanganesemia and neurodegeneration (Tuschl et al., 2016; Juneja et al., 2018; Marti-Sanchez et al., 2018; Rodan et al., 2018; Zeglam et al., 2018). In patients, subsequent accumulation of Mn in the globus pallidus, a component of the basal ganglia involved in motor control, leads to rapidly progressive dystonia-parkinsonism with onset in early childhood, a condition known as Hypermanganesemia with Dystonia 2 (HMNDYT2, OMIM # 617013). In a small number of patients, treatment has been attempted with Mn chelation using intravenous disodium calcium edetate (Na2CaEDTA) similar to a protocol established for HMNDYT1 (OMIM # 613280) caused by mutations in SLC30A10, a Mn exporter required for biliary excretion of Mn (Tuschl et al., 1993; Tuschl et al., 2012). MRI brain imaging of patients with either disorder are indistinguishable; hyperintensity of the basal ganglia, particularly the globus pallidus, and the white matter on T1-weighted imaging is a hallmark of both disorders (Tuschl et al., 2012; Tuschl et al., 2016). While patients with HMNDYT1 show significant improvement of neurological symptoms upon treatment initiation with stabilisation of the disease over many years (Tuschl et al., 2008; Tuschl et al., 2012), individuals with HMNDYT2 have variable treatment response, some even with worsening of their movement disorder (Tuschl et al., 2016; Marti-Sanchez et al., 2018). Consequently, the reasons for the difference in treatment response are poorly understood.

Although an essential trace metal, excess Mn has long been known to act as a neurotoxicant. Environmental Mn overexposure leads to preferential Mn accumulation in the globus pallidus similar to that observed in inherited Mn transporter defects, and causes manganism, a Parkinsonian movement disorder characterised by bradykinesia, akinetic rigidity, and dystonia, accompanied by psychiatric disturbances (Blanc, 2018; Chen et al., 2018). Despite its recognised role in neurodegenerative disease processes, we lack a deeper understanding of the mechanisms of Mn related neurotoxicity. The clinical similarities between manganism and Parkinson’s disease (PD) suggest that dopaminergic signalling is impaired upon Mn toxicity. However, in manganism, dopaminergic neurons within the substantia nigra are intact and response to L-DOPA is poor (Koller et al., 2004). Glutamatergic excitotoxicity as well as altered gamma-aminobutyric acid (GABA) signalling have also been proposed to underlie Mn associated neurodegeneration (Caito and Aschner, 2015). Indeed, Mn toxicity is likely mediated by a number of processes including oxidative stress, impaired mitochondrial function, protein misfolding and aggregation, and neuroinflammation (Martinez-Finley et al., 2013; Tjalkens et al., 2017).

We have recently established and characterised a zebrafish loss-of-function mutant *slc39a14^U801-/-^* that closely resembles the human phenotype with systemic accumulation of Mn, particularly in the brain (Tuschl et al., 2016). Homozygous mutants develop increased susceptibility to Mn toxicity and impaired locomotor behaviour upon Mn exposure. Mn levels can be lowered through chelation with Na_2_CaEDTA similar to what is observed in human patients (Troche et al., 2016).

In this study, we used RNA sequencing on individual larvae from an in-cross of heterozygous *slc39a14^U801^* zebrafish to identify novel potential targets of Mn toxicity. Furthermore, we determined the transcriptional signature elicited in response to MnCl_2_ treatment in mutant and sibling fish. Our results provide evidence that, in addition to Mn neurotoxicity, partial Mn deficiency that corrects upon Mn treatment is a prominent feature of *slc39a14* loss-of-function. We also determined that Ca^2+^ dyshomeostasis is a likely key event in both Mn deficiency and overload. Mn neurotoxicity appears to be further associated with activation of the unfolded protein response (UPR), oxidative stress, mitochondrial dysfunction, apoptosis and autophagy, and disruption of lysosomes and proteostasis.

## Materials and Methods

### Zebrafish husbandry

Zebrafish were reared on a 14/10 h light/dark cycle at 28.5°C. Embryos were obtained by natural spawning and staging was performed according to standard criteria (Kimmel et al., 1995). Previously generated *slc39a14^U801^* loss-of-function zebrafish and their siblings were used for all experiments (Tuschl et al., 2016). Ethical approval for zebrafish experiments was obtained from the Home Office UK under the Animal Scientific Procedures Act 1986.

### Preparation of larvae for RNA and DNA extraction

The progeny of a single in-cross of *slc39a14U801*^+/-^ fish were raised under standard conditions. At 2 dpf, the larvae were split into two groups and one group was exposed to MnCl_2_ added to the fishwater at a concentration of 50 μM. After 72 hours of exposure (at 5 dpf) single larvae were collected in the wells of a 96 well plate, immediately frozen on dry ice and stored at −80°C. For sequencing, frozen embryos were lysed in 100 μl RLT buffer (Qiagen) containing 1 μl of 14.3M beta mercaptoethanol (Sigma). The lysate was allowed to bind to 1.8 volumes of Agencourt RNAClean XP (Beckman Coulter) beads for 10 mins. The plate was then applied to a plate magnet (Invitrogen) until the solution cleared and the supernatant was removed without disturbing the beads. While still on the magnet the beads were washed three times with 70% ethanol and total nucleic acid was eluted from the beads as per the manufacturer’s instructions. Nucleic acid samples were used for genotyping of individual larvae by KASP assay (LGC Genomics) according to the manufacturer’s instructions and the following primers: wild-type allele 5’ GGCACATAATAATCCTCCATGGG 3’, mutant allele 5’ GGGCACATAATAATCCTCCATGGT 3’ and common primer 5’ CCCTGTATGTAGGCCTTCGGGTT 3’. After DNAse treatment, RNA was quantified using either Qubit RNA HS assay or Quant-iT RNA assay (Invitrogen).

### Transcript counting

DeTCT libraries were generated as described previously (Collins et al., 2015). Briefly, 300 ng of RNA from each genotyped sample was fragmented and bound to streptavidin beads. The 3’ ends of the fragmented RNA were pulled down using a biotinylated polyT primer. An RNA oligo containing the partial Illumina adapter 2 was ligated to the 5’ end of the bound fragment. The RNA fragment was eluted and reverse transcribed using an anchored oligo dT reverse transcriptase primer containing one of the 96 unique index sequences and part of the Illumina adapter 1. The Illumina adapters were completed during a library amplification step and the libraries were quantified using either the BioPhotometer (Eppendorf) or Pherastar (BMG Labtech). This was followed by size selection for an insert size of 70–270 bases. Equal quantities of libraries for each experiment were pooled, quantified by qPCR and sequenced on either HiSeq 2000 or HiSeq 2500.

Sequencing data were analysed as described previously (Collins et al., 2015). Briefly, sequencing reads were processed with the DeTCT detag_fastq.pl (https://github.com/iansealy/DETCT) script and aligned to the GRCz11 reference genome with BWA 0.5.10 (Li and Durbin, 2009). The resulting BAM files were processed using the DeTCT pipeline, which results in a list of regions (for simplicity referred to as genes in the Results) representing 3’ ends, together with a count for each sample. These counts were used for differential expression analysis with DESeq2 (Love et al., 2014). Each region was associated with Ensembl 95 (Yates et al., 2020) gene annotation based on the nearest transcript in the appropriate orientation. False positive 3’ ends, representing, for example, polyA-rich regions of the genome, were filtered using the DeTCT filter_output.pl script with the—strict option. Gene sets were analysed using the Cytoscape plugin ClueGO (Bindea et al., 2009) for gene ontology (GO) enrichment and Ontologizer (Bauer et al., 2008) for Zebrafish Anatomy Ontology (ZFA) enrichment.

### Quantitative real time PCR (qRT-PCR)

RNA extraction from 30 zebrafish larvae from the same genotype (homozygous mutant or wild-type) was performed using the TRIzol reagent (Invitrogen) according to the recommended protocol. DNA extraction was performed using the HotSHOT method (Truett et al., 2000). qRT-PCR was performed using GoTaq qPCR Master Mix (Promega) according to the recommended protocol. All samples were run in triplicates. qRT-PCR was carried out on a CFX96 Touch Real-Time PCR Detection System (BioRad). Only primer pairs with R2 values >0.99 and amplification efficiencies between 95% and 105% were used. Relative quantification of gene expression was determined using the 2^−ΔΔCt^ method, with elongation factor 1α (*ef1α*) as a reference gene (Livak and Schmittgen, 2001). The following primer sequences were used: *ef1α* forward 5’GTACTTCTCAGGCTGACTGTG3’, reverse 5’ ACGATCAGCTGTTTCACTCC3’; *bdnf* forward 5’AGATCGGCTGGCGGTTTATA3’, reverse 5’CATTGTGTACACTATCTGCCCC3’; *gnat2* forward 5’GCTGGCAGACGTCATCAAAA3’, reverse 5’CTCGGTGGGAAGGTAGTCAG3’; *hspa5* forward 5’GCTGGGCTGAATGTCATGAG3’, reverse 5’CAGCAGAGACACGTCAAAGG3’; *opn1mw2* forward 5’GCTGTCATTTCTGCGTTCCT3’, reverse 5’GACCATGCGTGTTACTTCCC3’; *pde6h* forward 5’CTCGCACCTTCAAGAGCAAG3’, reverse 5’CATGTCTCCAAACGCTTCCC3’; *prph2b* forward 5’GCCCTGGTGTCCTACTATGG3’, reverse 5’CTCTCGGGATTCTCTGGGTC3’.

### Optokinetic response (OKR)

The OKR was examined using a custom-built rig to track horizontal eye movements in response to whole-field motion stimuli. Larvae at 4 dpf were immobilised in 1.5% agarose in a 35 mm petri dish and analysed at 5 dpf. The agarose surrounding the eyes was removed to allow normal eye movements. Sinusoidal gratings with spatial frequencies of 0.05, 0.1, 0.13 and 0.16 cycles/degree were presented on a cylindrical diffusive screen 25 mm from the centre of the fish’s head. Gratings had a constant velocity of 10 degrees/second and changed direction and/or spatial frequency every 20 seconds. Eye movements were tracked under infrared illumination (720 nm) at 60 Hz using a Flea3 USB machine vision camera and custom-written software. A custom-designed Matlab code was used to determine the eye velocity (degrees per second).

### Retinal histology

5dpf larvae were fixed in 4% PFA overnight at 4°C. Dehydration was achieved by a series of increasing ethanol concentrations in PBS (50%, 70%, 80%, 90%, 95% and 100% ethanol). After dehydration larvae were incubated in a 1:1 ethanol Technovit 7100 solution (1% Hardener 1 in Technovit 7100 basic solution) for 1 h followed by incubation in 100% Technovit solution overnight at room temperature (Heraeus Kulzer, Germany). Larvae were than embedded in plastic moulds in Technovit 7100 polymerization medium and dried at 37°C for 1 h. Sections of 3 μm thickness were prepared with a microtome, mounted onto glass slides, and dried at 60°C. Sections were stained with Richardson (Romeis) solution (0.5% Borax, 0.5% Azur II, 0.5% Methylene Blue) and slides were mounted with Entellan (Merck, Darmstadt, Germany). Images were taken in the brightfield mode of a BX61 microscope (Olympus).

### Experimental design and statistical analyses

Animals were divided into four experimental groups: unexposed homozygous *slc39a14^U801-/-^* mutants and their siblings (wild-type and heterozygous genotypes), and MnCl_2_ exposed homozygous *slc39a14^U801-/-^* mutants and their siblings (wild-type and heterozygous genotypes). For the DeTCT data, an equal number of wild-type and heterozygous embryos were selected (see Fig. 1 for numbers of embryos for each experimental group). Embryos were all derived from a single cross to minimise the amount of biological variance not caused by the experimental conditions (i.e. genotype and Mn exposure). One wild-type Mn-exposed embryo was excluded from the data after visual inspection of the Principal Component Analysis as it did not group with any of the other samples. DESeq2 was used for differential expression analysis with the following model: ~ genotype + treatment + genotype:treatment. This models the observed counts as a function of the genotype (homozygous vs siblings) and the treatment (Mn exposed vs unexposed) and an interaction between the two and tests for significant parameters using the Wald test with a p value threshold of 0.05. For qRT-PCR and OKR analysis ANOVA with Tukey post-hoc testing was used to determine statistical significance, using the GraphPad Prism software (version 5). For GO term analysis, the settings for ClueGO were as follows: a right-sided hypergeometric test (enrichment only) was used with the Bonferroni step-down (Holm-Bonferroni) correction for multiple testing and terms with corrected p values >0.05 were discarded. For ZFA enrichment analysis, the Ontologizer Parent-Child-Union calculation method was used with Bonferroni correction.

**Fig. 1.**
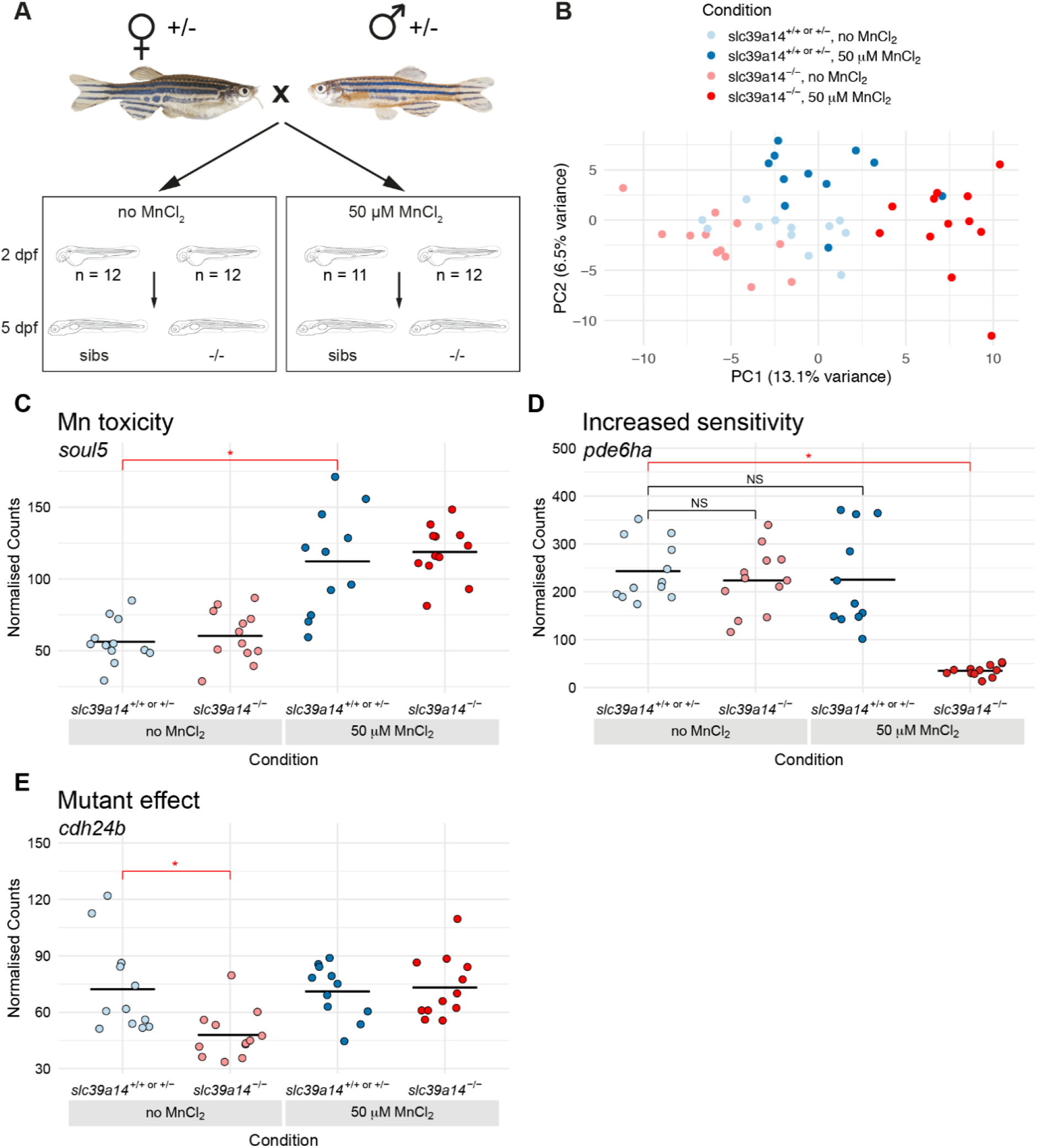
DeTCT analysis identifies three groups of differentially expressed genes. (A) Diagram of the experiment. Embryos from a *slc39a14^U801^* heterozygous in-cross were either exposed to 50 μM MnCl_2_ or left unexposed from 2 to 5 dpf. (B) Principal Component Analysis of the samples. Principal component (PC) 1 is plotted on the x-axis and PC2 on the y-axis. Samples belonging to the same condition group together. Unexposed sibling embryos are light blue and MnCl_2_ exposed ones are dark blue. Unexposed mutants are coloured light red and exposed mutants are dark red. (C) Group 1 (Mn toxicity) genes are defined as those with a significant difference between exposed and unexposed siblings (red bar with asterisk). Example plot of normalised counts for the *soul5* gene. The colour scheme for C–E is the same as in (B). (D) Group 2 (Increased sensitivity) genes are defined as those with a significant difference between exposed mutants and unexposed siblings (red bar with asterisk) without significant differences in either unexposed mutants or exposed siblings when compared to unexposed siblings (black bars labelled NS). (E) Group3 (Mutant effect) is defined as genes with a significant difference between unexposed mutants and unexposed siblings (red bar with asterisk).

### Transcription factor motif analysis

Transcription factor motif enrichment was performed using HOMER’s findMotifs.pl tool (v4.10.3) with default settings (Heinz et al., 2010). The GRCz11 promoter set used was created with HOMER’s updatePromoters.pl tool based on RefSeq genes from −2000 bp to 2000 bp relative to the TSS.

## Results

### Transcriptome analysis of *slc39a14^U801^* mutants identifies increased sensitivity to Mn toxicity and highlights additional Mn deficiency effects in homozygous mutants

To investigate the transcriptional profiles of *slc39a14^U801-/-^* mutants in the absence and presence of Mn treatment, embryos from a heterozygous in-cross were split into two groups and either raised under standard conditions (later referred to as unexposed), or treated with 50 μM MnCl_2_ from 2 until 5 days post fertilisation (dpf) (Fig. 1A). We have previously shown that this concentration elicits a locomotor phenotype in homozygous mutant larvae that is absent in siblings (Tuschl et al., 2016). We then carried out transcriptional profiling of individual 5 dpf larvae using 3’ tag sequencing (Collins et al., 2015). Principal Component Analysis (PCA) shows that the samples cluster according to genotype and treatment status (Fig. 1B). Analysis of differentially expressed genes between the four conditions produced three large sets of genes where each set had a characteristic expression profile. The first set are genes that are differentially expressed in MnCl_2_ exposed siblings compared with unexposed siblings (Fig. 1C, Mn toxicity) and represent a response to an increased concentration of Mn in the embryos. The second set contains genes that show increased sensitivity to Mn in *slc39a14^U801-/-^* mutants. These are defined as genes that are differentially expressed in MnCl_2_ exposed mutants compared with unexposed siblings, but not differentially expressed in unexposed mutants compared to unexposed siblings or exposed siblings compared with unexposed siblings (Fig. 1D, Increased sensitivity). The third set is composed of genes that are differentially expressed in unexposed mutants compared with unexposed siblings (Fig. 1E, Mutant effect). We will now consider these three groups of genes in turn (Table 1).

**Table 1.**
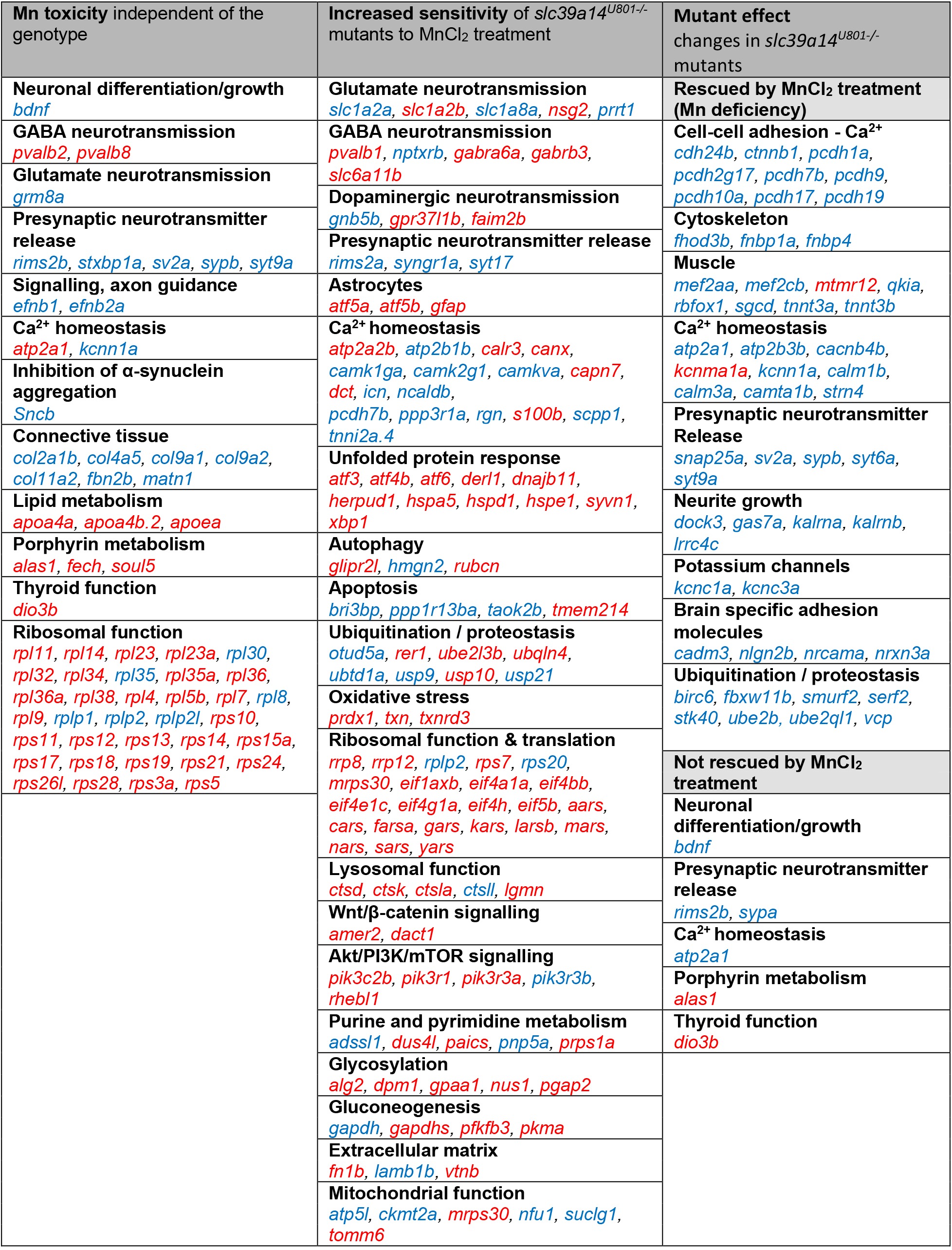
Differentially expressed genes grouped by function. Full lists in Supplementary Table 1. Red, increased gene expression. Blue, reduced gene expression.

### Mn toxicity causes differential gene expression independent from the genotype

MnCl_2_ treatment caused differential expression of 328 genes independent of the genotype (comparing MnCl_2_ exposed siblings and unexposed siblings) (Fig. 2A, Table 1 and Supplementary Table 1). Among them is brain-derived neurotrophic factor (*bdnf*) encoding a protein that is known to be altered upon Mn exposure (Zou et al., 2014). In addition, *bdnf* expression is also diminished in untreated mutants compared to siblings (Fig. 2B). Given that *slc39a14^U801-/-^* mutants show evidence of Mn toxicity already at 5 dpf (increased total Mn and reduced locomotor activity), this suggests that *bdnf* expression is a sensitive read-out for Mn toxicity. Mn associated suppression of BDNF signalling has been linked to diminished numbers of parvalbumin positive cells, mainly GABAergic interneurons (Fairless et al., 2019). Indeed, we find Parvalbumin encoding genes differentially expressed upon Mn exposure in mutants as well as siblings (*pvalb2, pvalb8*) and in treated mutants only (*pvalb1*). However, parvalbumin mRNA expression is upregulated in response to Mn in mutants and siblings, which is unexpected given the previously reported link to reduced numbers of Parvalbumin positive cells.

**Fig. 2.**
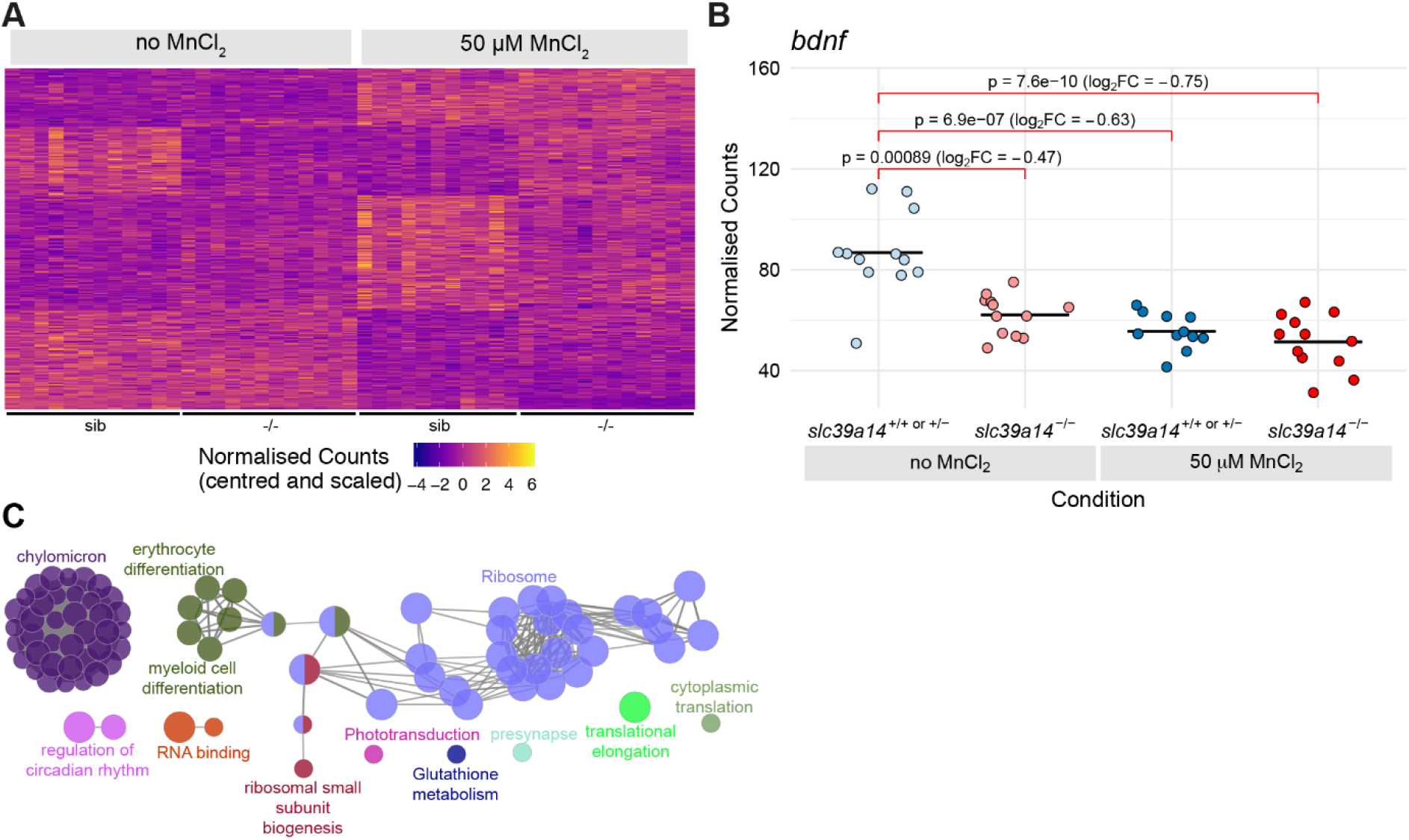
Manganese overexposure causes neurotoxicity and metabolic defects in wild-type zebrafish. (A) Heatmap of the expression of all 328 genes with a significant difference between exposed and unexposed siblings (Group 1 - Mn toxicity, Supplementary Table 1). Each row represents a different gene and each column is a sample. Mutant embryos are displayed for completeness although the group of genes is defined by the response in siblings only. The normalised counts for each gene have been mean centred and scaled by dividing by the standard deviation. (B) Plot of the normalised counts for each sample of a gene *(bdnf)* in Group 1. Unexposed sibling embryos are light blue and MnCl_2_ exposed ones are dark blue. Unexposed mutants are coloured light red and exposed mutants are dark red. (C) Enrichment of Gene Ontology (GO) terms associated with the genes in (A). Diagram produced using the CytoScape ClueGO App. Nodes represent enriched GO terms and edges connect GO terms that have annotated genes in common. Different components of the network are coloured according to the categories labelled on the diagram.

Among other brain-expressed genes affected by MnCl_2_ exposure in siblings are some involved in synaptic vesicle function (*rims2b, stxbp1a, sv2a, sypb, syt9a*), and genes encoding the metabotropic glutamate receptor (*grm8a*), β-synuclein (*sncb*) and ephrin-B membrane proteins (*efnb1, efnb2a*). Reduced ephrin-B levels have been linked to the pathophysiology of Alzheimer’s disease (AD) (Mroczko et al., 2018).

Mn is important for connective tissue integrity and bone mineralisation as a constituent of metalloenzymes and an enzyme co-factor (Sirri et al., 2016; Zofkova et al., 2017). Accordingly, our transcriptome analysis confirms that Mn exposure in zebrafish leads to reduced expression of multiple connective tissue related genes *(col2a1b, col4a5, col9a1a, col9a2, col11a2*, *dcn*, *fbn2b*, *matn1*).

Analysis of annotations to Gene Ontology (GO) terms shows enrichments of terms related to lipid metabolism (*apoa4b.2, apoa4a, apoea*), blood cell development *(alas1, fech, soul5;* Fig. 1C) and translation (35 ribosomal protein encoding genes) (Fig. 2C; Supplementary Table 2). Mn has previously been shown to interfere with heme-enzyme biogenesis and protein synthesis (Kaur et al., 2017; Chino et al., 2018; Hernandez et al., 2019).

### *slc39a14^U801-/-^* mutants show increased sensitivity to MnCl_2_ treatment compared to siblings

Our analysis showed that 613 genes are differentially expressed in MnCl_2_ exposed mutants compared with unexposed siblings, with no significant expression changes in either unexposed mutants or exposed siblings. Therefore, these are genes that show increased sensitivity to MnCl_2_ exposure in *slc39a14^U801-/-^* mutant larvae (Fig. 3A). 15% (95/613) of these genes also have a significant genotype-treatment interaction effect meaning that there is a synergistic effect on expression of treating mutant embryos with MnCl_2_ – i.e. the combined estimated effects of genotype and MnCl_2_ treatment alone are significantly less than the estimated log2 fold change for MnCl_2_ exposed mutants when compared to unexposed siblings (Fig. 3B, Table 1 and Supplementary Table 1). The remaining genes (518/613) show expression changes consistent with additive effects of the sub-significance threshold responses to genotype and MnCl_2_ exposure alone (Fig. 3C). Results from the transcriptome analysis were validated by qRT-PCR for a subset of six genes (*bdnf, gnat2, hspa5, opn1mw2, pde6h, prph2b*) using RNA extracted from equivalent embryos in a different experiment (Fig. 3D–E, Supplementary Fig. 1 and Supplementary Table 3). Changes in gene expression observed by qRT-PCR for all six genes were consistent with the results obtained from transcript counting (compare, for instance, Fig. 1D with Fig. 3E and Fig. 3B with Fig. 3D).

**Fig. 3.**
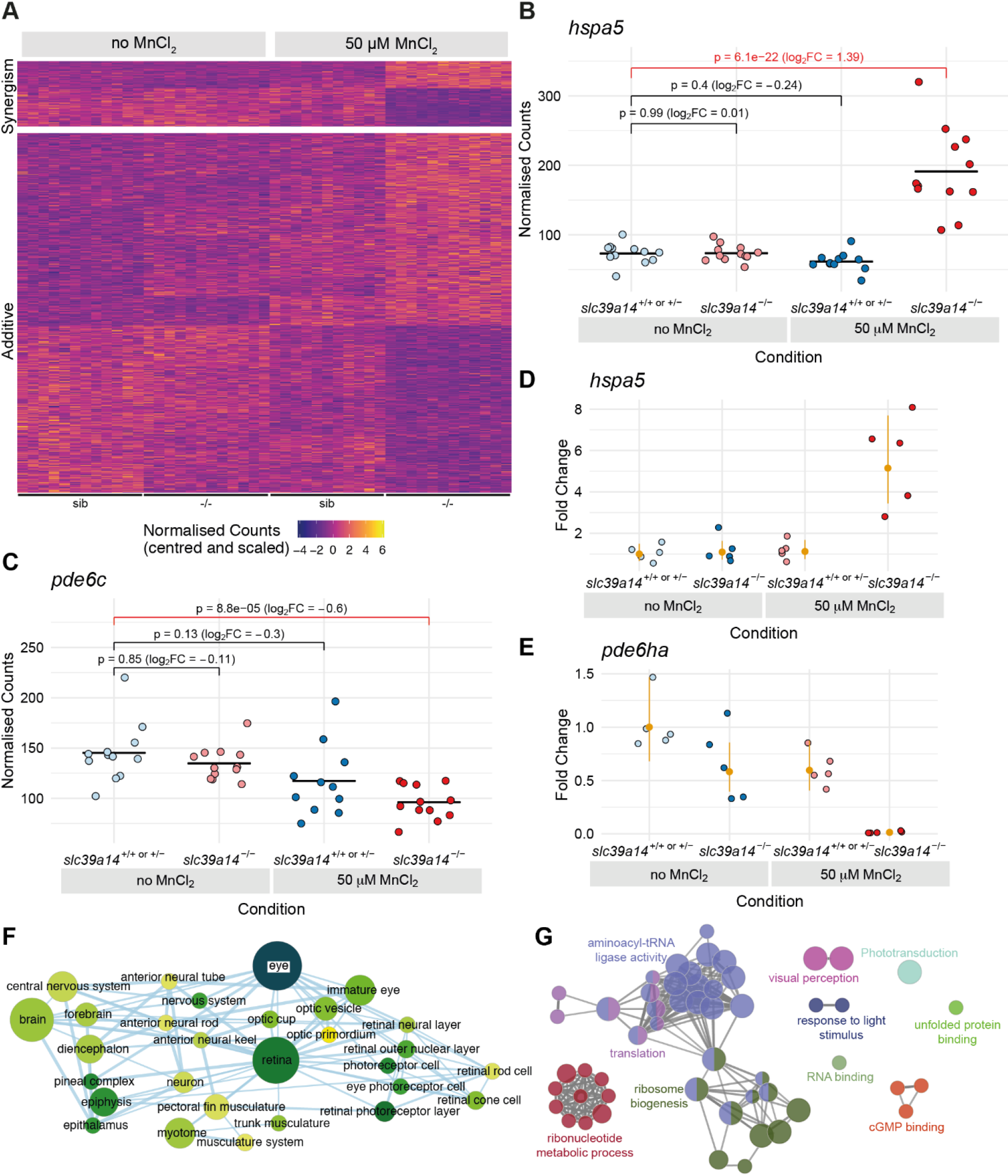
Effect of Mn treatment in *slc39a14^U801-/-^* mutants. (A) Heatmap of the expression of all genes (613) with a significant difference between exposed mutant and unexposed sibling embryos without significant treatment or genotype effects. The heatmaps are split into genes that show either synergistic or additive effects of the individual genotype and treatment effects. Each row represents a different gene and each column is a sample. The normalised counts for each gene have been mean centred and scaled by dividing by the standard deviation. (B) Example of a gene (*hspa5*) with a synergistic effect of treatment and genotype. The difference between the exposed mutants and unexposed siblings cannot be explained by adding together the separate effects of Mn treatment and the *slc39a14^U801^* mutation. Unexposed sibling embryos are light blue and MnCl_2_ exposed ones are dark blue. Unexposed mutants are coloured light red and exposed mutants are dark red. (C) Example of a gene (*pde6c*) that has an additive effect of treatment and genotype. The two sub-threshold effects of treatment and genotype produce the difference between exposed mutants and unexposed siblings when added together. Colour scheme as in (B). (D-E) qRT-PCR shows comparable gene expression changes as for the single embryo sequencing dataset. The individual samples are displayed as fold change relative to the mean value for unexposed siblings and the mean and 95% confidence intervals for each condition are in orange. (D) *hspa5*. Compare with (B). (E) *pde6ha*. Compare with Fig. 1D. (F) Enrichment Map network of the Zebrafish Anatomy Ontology (ZFA) enrichment results. Each node represents an enriched term and the edges join nodes that have overlapping genes annotated to them. The width of each edge is proportional to amount of overlap, nodes are coloured by -log_10_[Adjusted p value] and the size represents the number of significant genes annotated to the term. (G) ClueGO network diagram of the enrichment of Gene Ontology (GO) terms. Nodes represent enriched GO terms and edges connect nodes that share annotations to the significant genes. Different components of the network are coloured according to the categories as labelled on the diagram.

Enrichment of zebrafish anatomy (ZFA) terms shows that genes differentially expressed upon MnCl_2_ exposure in *slc39a14^U801-/-^* mutants are disproportionately expressed in the eye and nervous system (Fig. 3F; Supplementary Table 4). This is confirmed by the enrichment of GO terms such as visual perception and phototransduction. Also enriched are terms related to the ribosome, translation and the unfolded protein response (UPR) suggesting effects on protein production and folding (Fig. 3G and Supplementary Table 2).

### Increased sensitivity of *slc39a14^U801-/-^* mutants to MnCl_2_ treatment leads to Mn neurotoxicity

Enriched ZFA terms identified in MnCl_2_ exposed *slc39a14^U801-/-^* mutants that are not present in siblings confirm a high number of differentially expressed genes in the nervous system (Supplementary Table 4) consistent with the known role of Mn in neurotoxicity. Differentially expressed genes include several related to glutamatergic, GABAergic and dopaminergic signalling similar to previous studies that demonstrate impaired neurotransmitter signalling as a key event in Mn neurotoxicity (Marreilha Dos Santos et al., 2011). Genes with a link to glutamatergic circuitry include *slc1a2a* and *slc1a2b*, encoding the glutamate uptake transporter EAAT2, and *slc1a8a*, encoding a glutamate transporter present in teleosts only (Gesemann et al., 2010; Karki et al., 2015). Two genes required for the regulation of ionotropic AMPA type glutamate receptors (AMPAR) (*nsg2, prrt1*) show diminished expression in MnCl_2_ treated mutants (Chander et al., 2019; Troyano-Rodriguez et al., 2019).

Furthermore, we observe increased expression of *slc6a11b*, encoding a GABA uptake transporter, as well as the parvalbumin encoding gene (*pvalb1*) present in GABAergic interneurons. Expression of the GABA-A receptor encoding genes *gabra6a* and *gabrb3*, and *nptxrb*, encoding the neuronal pentraxin receptor expressed in parvalbumin positive interneurons (Kikuchihara et al., 2015), is reduced.

Differentially expressed genes associated with dopaminergic signalling include *gnb5b* and *gpr37l1b* that interact with neurotransmission via the D2 receptor, and *faim2b* for which loss-of-function increases susceptibility to dopaminergic neuron degeneration (Octeau et al., 2014; Komnig et al., 2016; Hertz et al., 2019). Furthermore, genes required for presynaptic neurotransmitter release (*rims2a, syngr1a, syt17*) show reduced expression. A role for astrocyte mediated Mn neurotoxicity and neuroinflammation is suggested by increased expression of the astrocyte related genes *atf5a*, *atf5b* and *gfap*.

### Increased sensitivity of *slc39a14^U801-/-^* mutants to MnCl_2_ treatment is associated with gene expression changes affecting calcium and protein homeostasis, and the unfolded protein response

Mn toxicity is known to cause protein misfolding and aggregation (Angeli et al., 2014; Harischandra et al., 2019b) and, as previously shown for Mn overexposure in *C. elegans* (Angeli et al., 2014), multiple genes involved in the UPR have increased expression in *slc39a14^U801-/-^* mutants upon MnCl_2_ treatment while siblings appear unaffected (Table 1 and Supplementary table 1). Ca^2+^ homeostasis within the endoplasmic reticulum (ER) plays a major role during the UPR and vice versa. Potentially linked to the UPR, over dozen of Ca^2+^associated/dependent genes are differentially expressed (Table 1). In MnCl_2_ treated *slc39a14^U801-/-^* mutants we observe differential expression of the Ca^2+^ ATPase encoding genes *atp2a2b* (SERCA2) and *atp2b1b* (PMCA1) as well as increased expression of genes encoding the Ca^2+^ chaperones calreticulin 3 (*calr3*) and calnexin (*canx*). Activation of the UPR as well as Ca^2+^ dyshomeostasis can promote apoptosis and autophagy. Concordantly, genes involved in autophagy and apoptosis are differentially expressed (Table 1). Degradation of misfolded and aggregated proteins occurs via the ubiquitin-proteasome system within the cytosol (Tamas et al., 2014) and MnCl_2_ exposed *slc39a14^U801-/-^* mutants show gene expression changes linked to ubiquitination (Table 1).

Oxidative stress and mitochondrial dysfunction are prominent features of Mn toxicity (Smith et al., 2017; Harischandra et al., 2019a). Consistent with this observation, essential genes of the thioredoxin/peroxiredoxin system *(prdx1, txn, txnrd3)* are activated in MnCl_2_ exposed *slc39a14^U801-/-^* mutants. Likewise, genes related to mitochondrial function show differential expression in MnCl_2_ treated mutants (Table 1).

As suggested by GO analysis, we observed pronounced expression changes of genes associated with ribosomal function and translation. MnCl_2_ treatment of *slc39a14^U801-/-^* mutants led to differential expression of eleven genes encoding tRNA synthetases, seven genes encoding translation initiation factors and six genes encoding ribosomal proteins. As mentioned above, Mn toxicity independent of the genotype led to differential expression of additional 33 ribosomal protein encoding genes suggesting that protein synthesis is a prominent target of Mn toxicity.

### Increased sensitivity of *slc39a14^U801-/-^* mutants to MnCl_2_ treatment manifests as impaired vision

30 genes involved in phototransduction were differentially expressed in MnCl_2_ exposed mutants but not in siblings (Fig. 4A, Supplementary Table 1). Hence, we further examined the vision of *slc39a14^U801-/-^* mutants. Raising *slc39a14^U801-/-^* mutant embryos/larvae on a 14 hour light, 10 hour dark cycle revealed absent visual background adaptation upon MnCl_2_ exposure while exposed wild-type larvae and unexposed mutants showed normal pigmentation (Fig. 4B). Visual background adaptation requires normal vision and is therefore impaired in blind larvae (Le et al., 2012). To determine whether *slc39a14^U801-/-^* larvae develop visual impairment, the optokinetic response (OKR) was analysed in homozygous *slc39a14^U801-/-^* larvae at 5 dpf after MnCl_2_ exposure. Exposed mutant larvae demonstrated a significant reduction in slow phase eye velocity at high spatial frequencies (Fig. 4C). Therefore, as predicted from the observed gene expression changes, Mn exposure leads to visual impairment and subsequent diminished visual background adaptation. Retinal histology appeared normal suggesting functional rather than overt structural deficits (Fig. 4D).

**Fig. 4.**
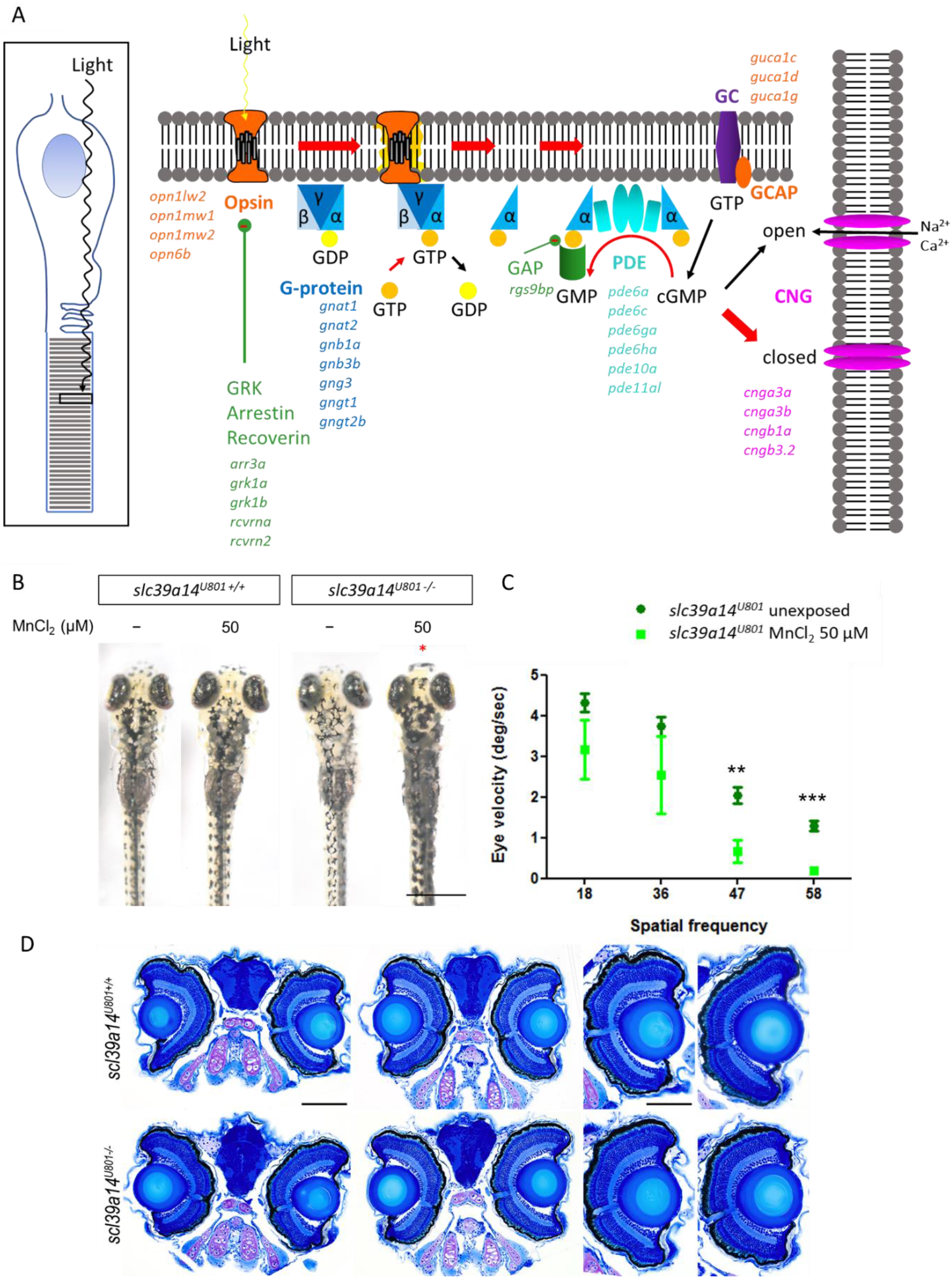
*slc39a14^u801^* loss-of-function mutants develop a visual phenotype upon MnCl_2_ expsoure. (A) Schematic showing the process of phototransduction (Kaupp and Seifert, 2002) with differentially expressed genes observed in MnCl_2_ exposed *slc39a14^U801-/-^* mutants in italics. cGMP, cyclic guanosine monophosphate. CNG, cyclic nucleotide gated non-selective cation channels. GC, guanylyl cyclase. GCAP, guanylate cyclase activating protein. PDE, phosphodiesterase. GRK, G-protein coupled receptor kinase. GAP, GTPase activating protein. (B) Dorsal views of wild-type siblings (*slc39a14*^*U801*+/+^, on the left) and *slc39a14^U801-/-^* larvae (on the right) at 5 dpf unexposed and exposed to 50 μM MnCl_2_. * indicates abnormal visual background adaptation. Scale bar 500 μm. (C) Graph showing the OKR (average of both eyes) of *slc39a14^U801-/-^* larvae unexposed (dark green squares) and exposed to 50 μM MnCl_2_ (light green circles). Data are presented as mean ± s.e.m. from five independent experiments. (**p<0.01; *** p<0.001). (D) Histologic analysis of retinal sections stained with Richardson–Romeis of wild-type siblings (*slc39a14*^*U801*+/+^, top row) and *slc39a14^U801-/-^* larvae (bottom row) at 5 dpf exposed to 50 μM MnCl_2_.

### Most genes affected in unexposed *slc39a14^U801-/-^* mutants are rescued by Mn treatment suggesting Mn deficiency

When compared to unexposed siblings, 266 genes show significantly different expression due to the U801 mutation alone (unexposed mutants versus unexposed siblings) (Fig. 5A; Supplementary table 1). Expression of 12% of these genes (31/266) is also significantly different between MnCl_2_ exposed mutants and unexposed siblings (Fig. 5B). Seven of these genes overlap with those differentially expressed in siblings upon MnCl_2_ exposure suggesting that these genes are sensitive targets of Mn toxicity (*alas1, atp2a1, bdnf, crim1, dio3b, dip2ca, rims2b*). However, the majority (88%, 235/266) of differentially expressed genes in unexposed mutants are not significantly differentially expressed when comparing MnCl_2_ exposed mutants and unexposed siblings (Fig. 5C). This suggests that the U801 mutation creates Mn deficiency leading to gene expression changes that are rescued by MnCl_2_ treatment towards levels observed in unexposed siblings.

**Fig. 5.**
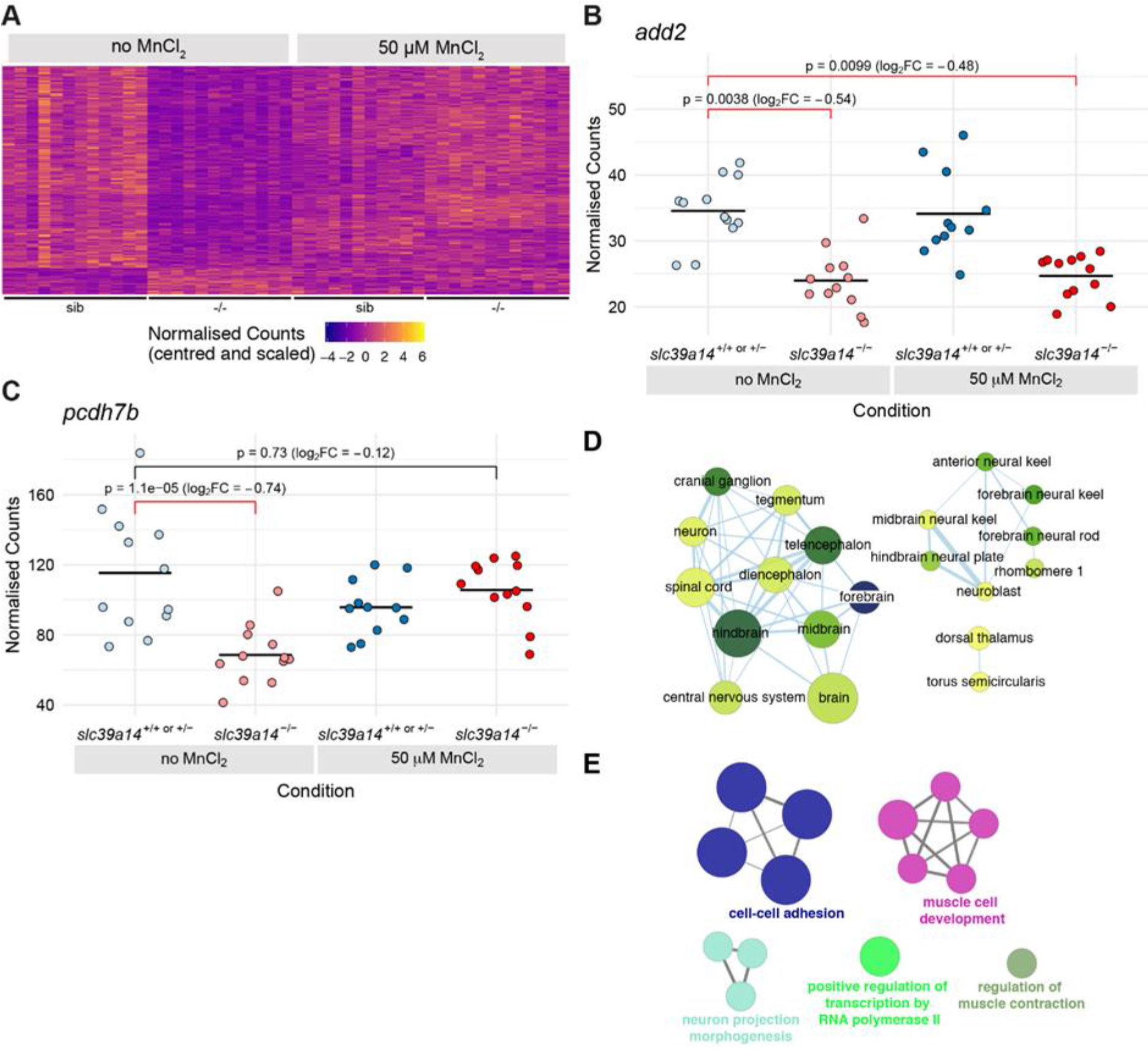
Exogenous Mn restores normal expression of many genes differentially expressed in unexposed *slc39a14^U801-/-^* mutants. (A) Heatmap of the expression of 266 genes with a significant difference between unexposed mutants and unexposed siblings. Each row represents a different gene and each column is a sample. The normalised counts for each gene have been mean centred and scaled by dividing by the standard deviation. (B) Plot of normalised counts for the *add2* gene. Expression is decreased in both unexposed and MnCl_2_ exposed mutant embryos. Unexposed sibling embryos are light blue and Mn-exposed ones are dark blue. Unexposed mutants are coloured light red and exposed mutants are dark red. (C) Plot of normalised counts for the *pcdh7b* gene. There are decreased counts in the unexposed mutant embryos that are rescued back to wild-type levels upon 50 μM MnCl_2_ treatment. Colour scheme as in (B). (C) Enrichment Map diagram of the enrichment of Zebrafish Anatomy Ontology (ZFA) terms for the genes differentially expressed in unexposed mutants that are rescued by Mn treatment. Nodes represent enriched ZFA terms and edges connect nodes that share annotations to the significant genes. The width of each edge is proportional to amount of overlap, nodes are coloured by -log_10_[Adjusted p value] and the size represents the number of significant genes annotated to the term. (D) ClueGO network diagram of the enrichment of Gene Ontology (GO) terms associated with the genes that are rescued by Mn treatment. Nodes represent enriched GO terms and edges connect nodes that share annotations to the significant genes. Different components of the network are coloured according to the categories as labelled on the diagram.

Zebrafish anatomy (ZFA) terms for the nervous system are enriched in this set of genes (Fig. 5D; Supplementary Table 4) and there is an enrichment for the GO terms cell-cell morphology, adhesion and cell-cell interactions (*cadm3, cdh24b, ctnnb1, fhod3b, fnbp1a, fnbp4, nlgn2b, nrcama, nrxn3a pcdh1a, pcdh2g17, pcdh7b, pcdh9, pcdh10a, pcdh17*) (Fig. 5E; Supplementary Table 2). Other brain expressed genes that change upon Mn deficiency include some essential for synaptic function and vesicle formation (*snap25a, sv2a, sypb, syt6a, syt9a*), neurite and axonal growth (*dock3, gas7a, kalrna, kalrnb, lrrc4c*) and potassium channels (*kcnc1a, kcnc3a*).

In addition, a group of differentially expressed Ca^2+^ associated genes are rescued by Mn treatment that is different to that observed upon Mn toxicity. These include genes encoding Ca^2+^ ATPases (*atp2a1*, *atp2b3b*), Ca^2+^ channels (*cacnb4b*), Ca^2+^ activated potassium channels (*kcnma1a, kcnn1a*), calmodulins (*calm1b, calm3a*) and calmodulin binding proteins (*camta1b, strn4*). Similarly, expression changes of genes involved in proteostasis and ubiquitination are observed in both Mn deficiency and toxicity, with a distinct affected gene set for each condition (Table 1).

### Both Mn toxicity and deficiency in *slc39a14^U801-/-^* mutants target the central nervous system

We analysed the three different gene sets for transcription factor motif enrichment using HOMER (Fig. 6A). The only enriched motifs we could identify were from the largest gene set identified in *slc39a14^U801-/-^* mutants upon MnCl_2_ treatment that were unchanged in treated siblings. The motifs included Chop/Atf4 which are part of the unfolded protein response (UPR), as well as HLF, NFIL3 and CEBP:AP1 (Supplementary Table 5). We next examined the enriched Zebrafish Anatomy Ontology (ZFA) terms for each gene set to identify tissue specificity of the observed gene expression changes (Fig. 6B). Whereas differentially expressed genes due to Mn toxicity effects independent of the genotype showed enrichment of ZFA terms primarily associated with liver and gut, the genes with differential expression due to Mn deficiency and increased sensitivity to Mn in *slc39a14^U801-/-^* mutants showed enrichment for the central nervous system (Supplementary Table 4).

**Fig. 6.**
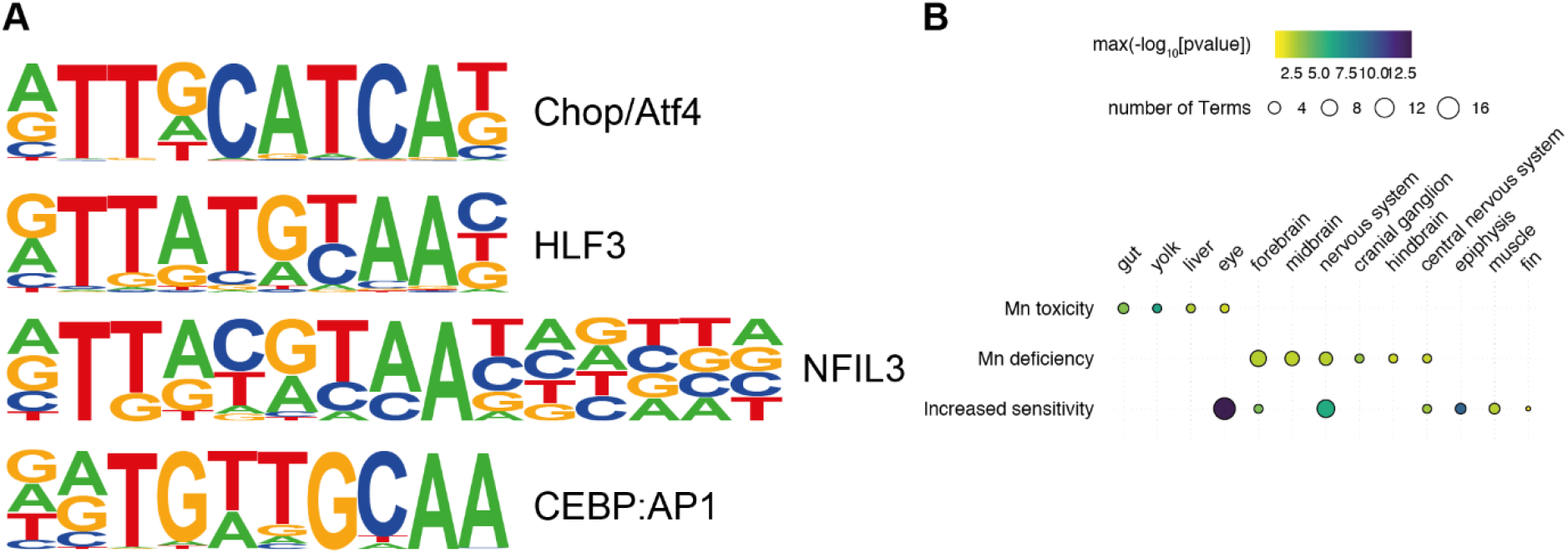
Comparative analysis of gene sets. (A) Example consensus binding motifs found to be enriched in the promoters of genes that show increased sensitivity to Mn treatment in *slc29a14^u801^* mutants (Group 1). The height of each base represents its frequency at that position in the consensus motif. (B) Bubble plot of the ZFA enrichment results across the three categories of response. Individual enriched ZFA terms were aggregated to the tissue/organ level. For example, the terms optic cup, retina and photoreceptor cell are all aggregated to the parent term eye. The size of each circle represents the number of individual terms enriched for the particular organ or tissue and they are coloured by the smallest of the p values (-log_10_ scaled).

## Discussion

Transcriptional profiling of *slc39a14^U801^* mutant zebrafish has identified distinct gene groups that are differentially expressed in normal physiological conditions and upon MnCl_2_ exposure. Consistent with the neurodegenerative phenotype observed in HMNDYT2 patients and the previously described accumulation of Mn in the brain of *slc39a14^U801-/-^* mutants (Tuschl et al., 2016), the majority of differentially expressed genes map to the CNS and the eye. Transcriptome analysis showed that Mn treatment leads to gene expression changes in both *slc39a14^U801-/-^* mutant and sibling zebrafish. Mutant larvae show differential expression of a much greater number of genes upon MnCl_2_ treatment that is not observed in treated siblings confirming an increased sensitivity to Mn toxicity. In addition, numerous differentially expressed genes in unexposed *slc39a14^U801-/-^* mutants normalised upon MnCl_2_ treatment. This suggests that Mn treatment in *slc39a14^U801-/-^* mutants rescues some of the transcriptomic changes observed in unexposed mutants. This implies that SLC39A14 loss leads to Mn deficiency in parallel to the observed Mn accumulation.

### Loss of *slc39a14* function in zebrafish causes Mn deficiency

Perhaps the most intriguing observation from the transcriptional profiling was that most differentially expressed genes in unexposed *slc39a14^U801-/-^* mutants normalised upon MnCl_2_ treatment. This indicates that whilst SLC39A14 deficiency leads to systemic Mn accumulation in some locations it also causes deficiency of Mn in some parts of the cell or specific types of cells due to its role as a Mn uptake transporter. One implication from this conclusion is that in patients, Mn chelation treatment would require careful monitoring in order to prevent over-chelation. Reducing Mn availability in parts of the cell may aggravate the neurological disease and lead to further decline. This partial Mn deficiency may explain why chelation therapy in patients with HMNDYT2 is less effective compared to those with HMNDYT1. There are only two individuals out of a dozen patients with HMNDYT2 reported in the literature who had a marked improvement upon Mn chelation (Tuschl et al., 2016; Rodan et al., 2018). Other treatment attempts have been less successful with some patients deteriorating upon Mn chelation (Tuschl et al., 2016; Marti-Sanchez et al., 2018).

The presence of Mn deficiency in *slc39a14^U801-/-^* mutants suggests that some features of HMNDYT2 may overlap with those observed in SLC39A8 deficiency, an inherited Mn transporter defect leading to systemic Mn deficiency (OMIM #616721). Affected individuals present with intellectual disability, developmental delay, hypotonia, epilepsy, strabismus, cerebellar atrophy and short stature (Boycott et al., 2015; Park et al., 2015). However, HMNDYT2 does not lead to any of these features aside from cerebellar atrophy described in some patients. SLC39A8 deficiency is also associated with dysglycosylation as Mn acts as a cofactor for the β-1,4-galactosyltransferase in the Golgi. However, transferrin glycosylation in HMNDYT2 is normal suggesting that Mn levels within the Golgi are not reduced (Tuschl et al., 2016).

The majority of differentially expressed genes in unexposed *slc39a14^U801-/-^* mutants that correct upon Mn treatment map to the CNS. As for Mn toxicity, several differentially expressed genes link to Ca^2+^ homeostasis and binding, however, these are different to those identified upon Mn overload. It is plausible that altered Mn levels, in both Mn deficiency and overload, result in Ca^2+^ dyshomeostasis. Expression of multiple genes encoding Ca^2+^ dependent cell-cell adhesion and interaction proteins, particularly protocadherins and formin related genes, is reduced in unexposed *slc39a14^U801-/-^* mutants. Protocadherins are mainly expressed in the CNS where they are required for normal neural circuitry activity and regulate synaptic function (Kim et al., 2011). Loss of protocadherin function in mice has been previously associated with neurodegeneration (Hasegawa et al., 2016). Formins are required for stabilisation of E-cadherins (Rao and Zaidel-Bar, 2016) which may link the changes observed in (proto-)cadherin expression with that of formin-associated genes. In addition, a number of genes required for Ca^2+^ triggered synaptic vesicle exocytosis was differentially expressed. Interestingly, synaptotagmin 1 can bind Ca^2+^ and Mn^2+^ in the same manner (Ubach et al., 1998). Mn dyshomeostasis may therefore directly affect neurotransmitter release.

### Unexposed *slc39a14^U801-/-^* mutants as well as MnCl_2_ treated mutants and siblings show evidence of Mn neurotoxicity

The mechanisms underlying Mn neurotoxicity are heterogenous suggesting an extensive role for Mn in brain pathobiology. Occupational manganism is associated with lower plasma BDNF levels (Zou et al., 2014), and Mn treatment in mice and rats reduces BDNF levels (Stansfield et al., 2014; Zhu et al., 2019). Indeed, *bdnf* expression is reduced in untreated *slc39a14^U801-/-^* mutants as well as MnCl_2_ exposed siblings confirming that *bdnf* expression is a sensitive readout of Mn neurotoxicity. BDNF promotes neuronal cell survival, neurite growth and cell migration, and as such is required for the postnatal growth of the striatum (Rauskolb et al., 2010).

In addition, Mn overexposure has previously been shown to disrupt neurotransmitter release via interaction with the SNARE complex which is mediated by increased intracellular Ca^2+^levels and subsequent activation of calpain, a Ca^2+^/Mn^2+^-activated neutral protease (Wang et al., 2018). Our results provide evidence that Mn neurotoxicity in *slc39a14^U801-/-^* mutants affects expression of genes encoding parts of the presynaptic neurotransmitter release machinery such as *rims2a, rims2b, syngr1a* and *syt17* as well as calpain (*capn7*).

The neuronal subtypes affected by Mn neurotoxicity remain subject of debate. Consistent with previous reports we observe altered expression of genes involved in glutamatergic, GABAergic and dopaminergic neurotransmission in MnCl_2_ treated *slc39a14^U801-/-^* mutants (Marreilha Dos Santos et al., 2017). Mn overexposure has been linked to impaired reuptake of glutamate from the synaptic cleft resulting in glutamatergic excitotoxicity (Erikson et al., 2008; Avila et al., 2010). In keeping with this finding, genes encoding glutamate uptake transporters (*slc1a2a, slc1a2b, slc1a8a*) as well as some required for the regulation of AMPA-type glutamate receptors (*nsg2, prrt1*) (Chander et al., 2019; Troyano-Rodriguez et al., 2019) are differentially expressed in MnCl_2_ exposed *slc39a14^U801-/-^* mutants. *SLC1A2* encodes the glutamate uptake transporter EAAT2 that is expressed on astrocytes and known to be downregulated upon MnCl_2_ exposure, subsequently leading to glutamate excitotoxicity (Karki et al., 2015).

In HMNDYT2 patients, Mn preferentially accumulates in the globus pallidus, a region that is particularly rich in GABAergic projections (Sidoryk-Wegrzynowicz and Aschner, 2013; Tuschl et al., 2016). In MnCl_2_ treated *slc39a14^U801-/-^* mutants expression of genes encoding the GABA-A receptor (*gabra6a, gabrb3*) and the GABA reuptake transporter (*slc6a11b*) is reduced. This is consistent with studies in rats where Mn exposure leads to diminished GABA-A receptor mRNA expression and interferes with GABA uptake in astrocytes (Fordahl and Erikson, 2014; Ou et al., 2017). Increased expression of genes encoding parvalbumin (*pvalb1, pbalb2* and *pvalb8*) in *slc39a14^U801-/-^* mutants and siblings upon MnCl_2_ treatment may be consistent with previous findings suggesting that GABAergic interneurons are a target of Mn neurotoxicity (Kikuchihara et al., 2015). Parvalbumin, a Ca^2+^ binding protein, can also bind Mn^2+^ with high affinity (Nara et al., 1994). Mn may therefore interact with parvalbumin directly or via changes in Ca^2+^ homeostasis. Mn exposure in mice leads to a reduction of parvalbumin positive cells likely due to suppression of BDNF signalling (Kikuchihara et al., 2015). Parvalbumin positive interneurons also express neuronal pentraxins and the neuronal pentraxin receptor. Pentraxins have previously been shown to play a role in neuroinflammation in PD and AD (Yin et al., 2009). Indeed, expression of *nptxrb* encoding the neuronal pentraxin receptor is reduced in *slc39a14^U801-/-^* mutants upon MnCl_2_ treatment.

Because manganism resembles Parkinson’s disease to some extent (both cause an akinetic movement disorder, albeit, with distinct clinical features) it seemed plausible that dopaminergic neurons are affected by Mn neurotoxicity. Indeed, several studies have shown dopaminergic neurodegeneration upon Mn exposure (Ijomone et al., 2016). However, transcriptome analysis of *slc39a14^U801-/-^* mutants identified changes in only three genes linked to dopaminergic signalling. *gnb5b* and *gpr37l1b* interact with neurotransmission via the D2 receptor, and loss-of-function of *faim2b* leads to increased susceptibility to dopaminergic neuron degeneration (Octeau et al., 2014; Komnig et al., 2016; Hertz et al., 2019). Therefore, it appears likely that interference with genes encoding proteins involved in dopaminergic circuitries is not the primary pathogenesis in *slc39a14^U801-/-^* mutants.

Neuroinflammation has been linked to Mn neurotoxicity supported by the observation that Mn predominantly accumulates in astrocytes rather than neurons (Tjalkens et al., 2017; Gorojod et al., 2018; Popichak et al., 2018). Indeed, Mn exposure in *slc39a14^U801^* loss-of-function mutants leads to differential expression of the astrocyte related genes *atf5a*, *atf5b* and *gfap*.

### Mn toxicity in *slc39a14^U801-/-^* mutants is associated with calcium dyshomeostasis, activation of the unfolded protein response and oxidative stress

Mn can replace Ca^2+^ in its biologically active sites and thereby affect Ca^2+^ homeostasis (Kalbitzer et al., 1978; Song et al., 2017). Mn overexposure increases intracellular Ca^2+^concentrations due to disruption of Ca^2+^ homeostasis at the mitochondria and the ER (Quintanar et al., 2012) and has previously been linked to neuronal loss and neurodegeneration (Choudhary et al., 2018; Ijomone et al., 2019). Chronically elevated Ca^2+^ levels leading to altered cellular signalling and mitochondrial damage is also a hallmark of neurodegeneration in PD (Ludtmann and Abramov, 2018). Indeed, Mn overload in *slc39a14^U801-/-^* mutants causes significant expression changes of Ca^2+^ associated genes. Impaired Ca^2+^ homeostasis may directly affect *bdnf* expression that is modulated by Ca^2+^/CaMK signalling (Liu et al., 2017). Ca^2+^ homeostasis is maintained by the ER, the key organelle in regulating proteostasis (Wang et al., 2012). ER stress is clearly evident in MnCl_2_ exposed *slc39a14^U801-/-^* mutants as multiple UPR associated genes are upregulated. HOMER analysis also confirms enrichment of the Chop/Atf4 motif in MnCL_2_ treated mutants. This is consistent with previous studies that show increased expression of ATF6 and HSPA5 as well as increased Xbp1 mRNA splicing in Mn exposed brain slices (Xu et al., 2013). ER stress increases the expression of calcium pumps and chaperones such as calreticulin which help to alleviate protein misfolding while dysfunctional Ca^2+^ chaperones cause activation of the UPR (Carreras-Sureda et al., 2018). Calreticulin and calnexin act together as a quality control system that causes retention of misfolded proteins within the ER (McCaffrey and Braakman, 2016). Expression of both genes is increased in MnCl_2_ exposed *slc39a14^U801-/-^* mutants.

Generation of reactive oxygen species (ROS) with subsequent oxidative stress and mitochondrial dysfunction is a hallmark of neurodegenerative disorders as well as metal toxicity and contributes to protein misfolding (Gomez and Germain, 2019; Harischandra et al., 2019a). The thioredoxin/peroxiredoxin system required for the reduction of H2O2 protects cells from oxidative stress (Samet and Wages, 2018). Oxidative stress is highlighted by the upregulation of the thioredoxin/thioredoxin reductase and peroxiredoxin system in MnCl_2_ exposed *slc39a14^U801-/-^* loss-of-function mutants, similar to previous reports in rats (Taka et al., 2012). Increased ROS generation itself can cause Ca^2+^ dyshomeostasis, lysosomal impairment, abnormal protein folding and mitochondrial dysfunction (Gorlach et al., 2015; Harischandra et al., 2019a). ROS leads to oxidation of the thiol group in cysteines of Ca^2+^ channels and pumps thereby affecting intracellular Ca^2+^ levels (Zhang et al., 2016). Furthermore, ROS cause apoptosis and autophagy via lysosomal membrane permeabilisation and cathepsin release (Gorojod et al., 2017; Wang et al., 2017; Porte Alcon et al., 2018; Zhi et al., 2019). Consistent with this observation, the key autophagy gene *rubcn*, encoding a beclin 1 interactor and responsible for autophagy initiation (Liu et al., 2019), is upregulated in *slc39a14^U801-/-^* mutants due to Mn overload. In addition, cathepsin gene expression is altered in MnCl_2_ treated *slc39a14^U801-/-^* mutants linking manganese to dysregulation of lysosomal function and autophagy as previously suggested (Zhang et al., 2019).

### Mn toxicity interferes with protein synthesis and metabolism

As suggested by GO term enrichment analysis, MnCl_2_ exposure led to differential expression of multiple genes encoding ribosomal proteins, tRNA synthetases and translation initiation factors in *slc39a14^U801-/-^* mutants. Interference of Mn with protein synthesis has been identified in yeast where Mn overexposure leads to reduced total rRNA levels and diminished ribosome formation (Hernandez et al., 2019). In addition, MnCl_2_ exposure in *slc39a14^U801-/-^* mutants is associated with gene expression changes linked to the Ubiquitination/Proteasome System (UPS). The UPS, essential for protein quality control, is susceptible to oxidative stress (Li et al., 2011; Zhang et al., 2016). Misregulation of the UPS has causally been linked to neurodegeneration in PD (Walden and Muqit, 2017). Heavy metals impair protein folding and promote protein aggregation suggesting that Mn can equally contribute to protein misfolding (Tamas et al., 2014).

### Mn toxicity in in *slc39a14^U801-/-^* zebrafish causes a visual phenotype

Interestingly, transcriptome analysis revealed an unsuspected Mn toxicity effect in *slc39a14^U801-/-^* zebrafish, a pronounced visual phenotype characterised by impaired visual background adaptation and impaired OKR upon MnCl_2_ exposure. To date, retinal Mn toxicity has not been previously reported in affected patients or animal models. Neither environmental overexposure nor systemic Mn accumulation in HMNDYT1 and HMNDYT2 lead to impaired vision in humans. Inherited Mn transporter defects have only recently been reported and it is possible that visual function becomes affected only in later life. Indeed, both Mn uptake transporters, SLC39A8 and SLC39A14, are highly expressed in the retinal pigment epithelium (RPE) (Leung et al., 2008). It has previously been shown that other heavy metals such as cadmium and lead accumulate in ocular tissues, particularly in the RPE (Erie et al., 2005). Mn plays an essential role in retinal function where it is required for normal ultrastructure of the retina (Gong and Amemiya, 1996). Possible differences between the human and zebrafish phenotype may simply be caused by the direct contact of the zebrafish eye with Mn in the fishwater contributing to enhanced ocular Mn uptake and toxicity.

In conclusion, our results demonstrate that partial Mn deficiency is an additional key feature of *slc39a14* deficiency in zebrafish which should be considered in the treatment of affected individuals with SLC39A14 mutations. The *slc39a14^U801^* loss-of-function zebrafish mutant is proving an invaluable disease model to study the disease pathogenesis of HMNDYT2 as well as Mn neurotoxicity in general.

## Acknowledgments

K.T. was supported by Action Medical Research (GN1999), the Academy of Medical Sciences, the National Institute for Health Research (NIHR, Academic Clinical Lectureship) and the Great Ormond Street Hospital Charity (V0018). K.T., S.C.F.N and S.W.W. were supported by the UCL Neuroscience ZNZ Collaboration. L.E.V. was funded by FONDECYT grant (11160951), CONICYT International network grants (REDI170300 and REDES170010), and Universidad Mayor FDP grant (PEP I-2019074). S.W. was supported by the MRC (MR/L003775/1) and Wellcome Trust (104682/Z/14/Z). E.B.N. was supported by core grants to the Wellcome Sanger Institute (WT098051 and 206194).

This publication presents independent research funded by the National Institute for Health Research (NIHR). The views expressed are those of the authors and not necessarily those of the NHS, the NIHR or the Department of Health and Social Care.

We thank Dr Philippa Mills and Prof Peter Clayton for their input and fruitful discussions. We are also grateful to Neha Wali and the Sanger Institute sequencing pipelines for sample processing and sequencing.

## References

Angeli S, Barhydt T, Jacobs R, Killilea DW, Lithgow GJ, Andersen JK (2014) Manganese disturbs metal and protein homeostasis in Caenorhabditis elegans. Metallomics 6:1816–1823.

Avila DS, Colle D, Gubert P, Palma AS, Puntel G, Manarin F, Noremberg S, Nascimento PC, Aschner M, Rocha JB, Soares FA (2010) A possible neuroprotective action of a vinylic telluride against Mn-induced neurotoxicity. Toxicol Sci 115:194–201.

Bauer S, Grossmann S, Vingron M, Robinson PN (2008) Ontologizer 2.0--a multifunctional tool for GO term enrichment analysis and data exploration. Bioinformatics 24:1650–1651.

Bindea G, Mlecnik B, Hackl H, Charoentong P, Tosolini M, Kirilovsky A, Fridman WH, Pages F, Trajanoski Z, Galon J (2009) ClueGO: a Cytoscape plug-in to decipher functionally grouped gene ontology and pathway annotation networks. Bioinformatics 25:1091–1093.

Blanc PD (2018) The early history of manganese and the recognition of its neurotoxicity, 1837-1936. Neurotoxicology 64:5–11.

Boycott KM et al. (2015) Autosomal-Recessive Intellectual Disability with Cerebellar Atrophy Syndrome Caused by Mutation of the Manganese and Zinc Transporter Gene SLC39A8. Am J Hum Genet 97:886–893.

Caito S, Aschner M (2015) Neurotoxicity of metals. Handb Clin Neurol 131:169–189.

Carreras-Sureda A, Pihan P, Hetz C (2018) Calcium signaling at the endoplasmic reticulum: fine-tuning stress responses. Cell Calcium 70:24–31.

Chander P, Kennedy MJ, Winckler B, Weick JP (2019) Neuron-Specific Gene 2 (NSG2) Encodes an AMPA Receptor Interacting Protein That Modulates Excitatory Neurotransmission. eNeuro 6.

Chen P, Bornhorst J, Aschner M (2018) Manganese metabolism in humans. Front Biosci (Landmark Ed) 23:1655–1679.

Chino M, Leone L, Zambrano G, Pirro F, D’Alonzo D, Firpo V, Aref D, Lista L, Maglio O, Nastri F, Lombardi A (2018) Oxidation catalysis by iron and manganese porphyrins within enzyme-like cages. Biopolymers 109:e23107.

Choudhary B, Mandelkow E, Mandelkow EM, Pir GJ (2018) Glutamatergic nervous system degeneration in a C. elegans Tau(A152T) tauopathy model involves pathways of excitotoxicity and Ca(2+) dysregulation. Neurobiol Dis 117:189–202.

Erie JC, Butz JA, Good JA, Erie EA, Burritt MF, Cameron JD (2005) Heavy metal concentrations in human eyes. Am J Ophthalmol 139:888–893.

Erikson KM, Dorman DC, Lash LH, Aschner M (2008) Duration of airborne-manganese exposure in rhesus monkeys is associated with brain regional changes in biomarkers of neurotoxicity. Neurotoxicology 29:377–385.

Fairless R, Williams SK, Diem R (2019) Calcium-Binding Proteins as Determinants of Central Nervous System Neuronal Vulnerability to Disease. Int J Mol Sci 20.

Fordahl SC, Erikson KM (2014) Manganese accumulation in membrane fractions of primary astrocytes is associated with decreased gamma-aminobutyric acid (GABA) uptake, and is exacerbated by oleic acid and palmitate. Environ Toxicol Pharmacol 37:1148–1156.

Gesemann M, Lesslauer A, Maurer CM, Schonthaler HB, Neuhauss SC (2010) Phylogenetic analysis of the vertebrate excitatory/neutral amino acid transporter (SLC1/EAAT) family reveals lineage specific subfamilies. BMC Evol Biol 10:117.

Gomez M, Germain D (2019) Cross talk between SOD1 and the mitochondrial UPR in cancer and neurodegeneration. Mol Cell Neurosci 98:12–18.

Gong H, Amemiya T (1996) Ultrastructure of retina of manganese-deficient rats. Invest Ophthalmol Vis Sci 37:1967–1974.

Gorlach A, Bertram K, Hudecova S, Krizanova O (2015) Calcium and ROS: A mutual interplay. Redox Biol 6:260–271.

Gorojod RM, Alaimo A, Porte Alcon S, Saravia F, Kotler ML (2017) Interplay between lysosomal, mitochondrial and death receptor pathways during manganese-induced apoptosis in glial cells. Arch Toxicol 91:3065–3078.

Gorojod RM, Alaimo A, Porte Alcon S, Martinez JH, Cortina ME, Vazquez ES, Kotler ML (2018) Heme Oxygenase-1 protects astroglia against manganese-induced oxidative injury by regulating mitochondrial quality control. Toxicol Lett 295:357–368.

Harischandra DS, Ghaisas S, Zenitsky G, Jin H, Kanthasamy A, Anantharam V, Kanthasamy AG (2019a) Manganese-Induced Neurotoxicity: New Insights Into the Triad of Protein Misfolding, Mitochondrial Impairment, and Neuroinflammation. Front Neurosci 13:654.

Harischandra DS, Rokad D, Neal ML, Ghaisas S, Manne S, Sarkar S, Panicker N, Zenitsky G, Jin H, Lewis M, Huang X, Anantharam V, Kanthasamy A, Kanthasamy AG (2019b) Manganese promotes the aggregation and prion-like cell-to-cell exosomal transmission of alpha-synuclein. Sci Signal 12.

Hasegawa S, Kumagai M, Hagihara M, Nishimaru H, Hirano K, Kaneko R, Okayama A, Hirayama T, Sanbo M, Hirabayashi M, Watanabe M, Hirabayashi T, Yagi T (2016) Distinct and Cooperative Functions for the Protocadherin-alpha, -beta and -gamma Clusters in Neuronal Survival and Axon Targeting. Front Mol Neurosci 9:155.

Heinz S, Benner C, Spann N, Bertolino E, Lin YC, Laslo P, Cheng JX, Murre C, Singh H, Glass CK (2010) Simple combinations of lineage-determining transcription factors prime cis-regulatory elements required for macrophage and B cell identities. Mol Cell 38:576–589.

Hernandez RB, Moteshareie H, Burnside D, McKay B, Golshani A (2019) Manganese-induced cellular disturbance in the baker’s yeast, Saccharomyces cerevisiae with putative implications in neuronal dysfunction. Sci Rep 9:6563.

Hertz E, Terenius L, Vukojevic V, Svenningsson P (2019) GPR37 and GPR37L1 differently interact with dopamine 2 receptors in live cells. Neuropharmacology 152:51–57.

Ijomone OM, Miah MR, Peres TV, Nwoha PU, Aschner M (2016) Null allele mutants of trt-1, the catalytic subunit of telomerase in Caenorhabditis elegans, are less sensitive to Mn-induced toxicity and DAergic degeneration. Neurotoxicology 57:54–60.

Ijomone OM, Aluko OM, Okoh COA, Martins AC, Jr., Aschner M (2019) Role for calcium signaling in manganese neurotoxicity. J Trace Elem Med Biol 56:146–155.

Juneja M, Shamim U, Joshi A, Mathur A, Uppili B, Sairam S, Ambawat S, Dixit R, Faruq M (2018) A novel mutation in SLC39A14 causing hypermanganesemia associated with infantile onset dystonia. J Gene Med 20:e3012.

Kalbitzer HR, Stehlik D, Hasselbach W (1978) The binding of calcium and magnesium to sarcoplasmic reticulum vesicles as studied by manganese electron paramagnetic resonance. Eur J Biochem 82:245–255.

Karki P, Smith K, Johnson J, Jr., Aschner M, Lee EY (2015) Genetic dys-regulation of astrocytic glutamate transporter EAAT2 and its implications in neurological disorders and manganese toxicity. Neurochem Res 40:380–388.

Kaupp UB, Seifert R (2002) Cyclic nucleotide-gated ion channels. Physiol Rev 82:769–824.

Kaur G, Kumar V, Arora A, Tomar A, Ashish, Sur R, Dutta D (2017) Affected energy metabolism under manganese stress governs cellular toxicity. Sci Rep 7:11645.

Kikuchihara Y, Abe H, Tanaka T, Kato M, Wang L, Ikarashi Y, Yoshida T, Shibutani M (2015) Relationship between brain accumulation of manganese and aberration of hippocampal adult neurogenesis after oral exposure to manganese chloride in mice. Toxicology 331:24–34.

Kim SY, Yasuda S, Tanaka H, Yamagata K, Kim H (2011) Non-clustered protocadherin. Cell Adh Migr 5:97–105.

Kimmel CB, Ballard WW, Kimmel SR, Ullmann B, Schilling TF (1995) Stages of embryonic development of the zebrafish. Dev Dyn 203:253–310.

Koller WC, Lyons KE, Truly W (2004) Effect of levodopa treatment for parkinsonism in welders: A double-blind study. Neurology 62:730–733.

Komnig D, Schulz JB, Reich A, Falkenburger BH (2016) Mice lacking Faim2 show increased cell death in the MPTP mouse model of Parkinson disease. J Neurochem 139:848–857.

Le HG, Dowling JE, Cameron DJ (2012) Early retinoic acid deprivation in developing zebrafish results in microphthalmia. Vis Neurosci 29:219–228.

Leung KW, Liu M, Xu X, Seiler MJ, Barnstable CJ, Tombran-Tink J (2008) Expression of ZnT and ZIP zinc transporters in the human RPE and their regulation by neurotrophic factors. Invest Ophthalmol Vis Sci 49:1221–1231.

Li H, Durbin R (2009) Fast and accurate short read alignment with Burrows-Wheeler transform. Bioinformatics 25:1754–1760.

Li H, Wu S, Shi N, Lian S, Lin W (2011) Nrf2/HO-1 pathway activation by manganese is associated with reactive oxygen species and ubiquitin-proteasome pathway, not MAPKs signaling. J Appl Toxicol 31:690–697.

Liu K, Guo C, Lao Y, Yang J, Chen F, Zhao Y, Yang Y, Yang J, Yi J (2019) A fine-tuning mechanism underlying self-control for autophagy: deSUMOylation of BECN1 by SENP3. Autophagy:1–16.

Liu SH, Lai YL, Chen BL, Yang FY (2017) Ultrasound Enhances the Expression of Brain-Derived Neurotrophic Factor in Astrocyte Through Activation of TrkB-Akt and Calcium-CaMK Signaling Pathways. Cereb Cortex 27:3152–3160.

Livak KJ, Schmittgen TD (2001) Analysis of relative gene expression data using real-time quantitative PCR and the 2(-Delta Delta C(T)) Method. Methods 25:402–408.

Love MI, Huber W, Anders S (2014) Moderated estimation of fold change and dispersion for RNA-seq data with DESeq2. Genome Biol 15:550.

Ludtmann MHR, Abramov AY (2018) Mitochondrial calcium imbalance in Parkinson’s disease. Neurosci Lett 663:86–90.

Marreilha Dos Santos AP, Andrade V, Aschner M (2017) Neuroprotective and Therapeutic Strategies for Manganese-Induced Neurotoxicity. Clin Pharmacol Transl Med 1:54–62.

Marreilha Dos Santos AP, Lopes SM, Batoreu MC, Aschner M (2011) Prolactin is a peripheral marker of manganese neurotoxicity. Brain Res 1382:282–290.

Marti-Sanchez L, Ortigoza-Escobar JD, Darling A, Villaronga M, Baide H, Molero-Luis M, Batllori M, Vanegas MI, Muchart J, Aquino L, Artuch R, Macaya A, Kurian MA, Duenas P (2018) Hypermanganesemia due to mutations in SLC39A14: further insights into Mn deposition in the central nervous system. Orphanet J Rare Dis 13:28.

Martinez-Finley EJ, Gavin CE, Aschner M, Gunter TE (2013) Manganese neurotoxicity and the role of reactive oxygen species. Free Radic Biol Med.

McCaffrey K, Braakman I (2016) Protein quality control at the endoplasmic reticulum. Essays Biochem 60:227–235.

Mroczko B, Groblewska M, Litman-Zawadzka A, Kornhuber J, Lewczuk P (2018) Cellular Receptors of Amyloid beta Oligomers (AbetaOs) in Alzheimer’s Disease. Int J Mol Sci 19.

Nara M, Tasumi M, Tanokura M, Hiraoki T, Yazawa M, Tsutsumi A (1994) Infrared studies of interaction between metal ions and Ca(2+)-binding proteins. Marker bands for identifying the types of coordination of the side-chain COO- groups to metal ions in pike parvalbumin (pI = 4.10). FEBS Lett 349:84–88.

Octeau JC, Schrader JM, Masuho I, Sharma M, Aiudi C, Chen CK, Kovoor A, Celver J (2014) G protein beta 5 is targeted to D2-dopamine receptor-containing biochemical compartments and blocks dopamine-dependent receptor internalization. PLoS One 9:e105791.

Ou CY, Luo YN, He SN, Deng XF, Luo HL, Yuan ZX, Meng HY, Mo YH, Li SJ, Jiang YM (2017) Sodium P-Aminosalicylic Acid Improved Manganese-Induced Learning and Memory Dysfunction via Restoring the Ultrastructural Alterations and gamma-Aminobutyric Acid Metabolism Imbalance in the Basal Ganglia. Biol Trace Elem Res 176:143–153.

Park JH et al. (2015) SLC39A8 Deficiency: A Disorder of Manganese Transport and Glycosylation. Am J Hum Genet 97:894–903.

Popichak KA, Afzali MF, Kirkley KS, Tjalkens RB (2018) Glial-neuronal signaling mechanisms underlying the neuroinflammatory effects of manganese. J Neuroinflammation 15:324.

Porte Alcon S, Gorojod RM, Kotler ML (2018) Regulated Necrosis Orchestrates Microglial Cell Death in Manganese-Induced Toxicity. Neuroscience 393:206–225.

Quintanar L, Montiel T, Marquez M, Gonzalez A, Massieu L (2012) Calpain activation is involved in acute manganese neurotoxicity in the rat striatum in vivo. Exp Neurol 233:182–192.

Rao MV, Zaidel-Bar R (2016) Formin-mediated actin polymerization at cell-cell junctions stabilizes E-cadherin and maintains monolayer integrity during wound repair. Mol Biol Cell 27:2844–2856.

Rauskolb S, Zagrebelsky M, Dreznjak A, Deogracias R, Matsumoto T, Wiese S, Erne B, Sendtner M, Schaeren-Wiemers N, Korte M, Barde YA (2010) Global deprivation of brain-derived neurotrophic factor in the CNS reveals an area-specific requirement for dendritic growth. J Neurosci 30:1739–1749.

Rodan LH, Hauptman M, D’Gama AM, Qualls AE, Cao S, Tuschl K, Al-Jasmi F, Hertecant J, Hayflick SJ, Wessling-Resnick M, Yang ET, Berry GT, Gropman A, Woolf AD, Agrawal PB (2018) Novel founder intronic variant in SLC39A14 in two families causing Manganism and potential treatment strategies. Mol Genet Metab 124:161–167.

Samet JM, Wages PA (2018) Oxidative Stress from Environmental Exposures. Curr Opin Toxicol 7:60–66.

Sidoryk-Wegrzynowicz M, Aschner M (2013) Manganese toxicity in the central nervous system: the glutamine/glutamate-gamma-aminobutyric acid cycle. J Intern Med 273:466–477.

Sirri F, Maiorano G, Tavaniello S, Chen J, Petracci M, Meluzzi A (2016) Effect of different levels of dietary zinc, manganese, and copper from organic or inorganic sources on performance, bacterial chondronecrosis, intramuscular collagen characteristics, and occurrence of meat quality defects of broiler chickens. Poult Sci 95:1813–1824.

Smith MR, Fernandes J, Go YM, Jones DP (2017) Redox dynamics of manganese as a mitochondrial life-death switch. Biochem Biophys Res Commun 482:388–398.

Song D, Ma J, Chen L, Guo C, Zhang Y, Chen T, Zhang S, Zhu Z, Tian L, Niu P (2017) FOXO3 promoted mitophagy via nuclear retention induced by manganese chloride in SH-SY5Y cells. Metallomics 9:1251–1259.

Stansfield KH, Bichell TJ, Bowman AB, Guilarte TR (2014) BDNF and Huntingtin protein modifications by manganese: implications for striatal medium spiny neuron pathology in manganese neurotoxicity. J Neurochem 131:655–666.

Taka E, Mazzio E, Soliman KF, Renee RR (2012) Microarray genomic profile of mitochondrial and oxidant response in manganese chloride treated PC12 cells. Neurotoxicology 33:162–168.

Tamas MJ, Sharma SK, Ibstedt S, Jacobson T, Christen P (2014) Heavy metals and metalloids as a cause for protein misfolding and aggregation. Biomolecules 4:252–267.

Thompson KJ, Wessling-Resnick M (2019) ZIP14 is degraded in response to manganese exposure. Biometals 32:829–843.

Tjalkens RB, Popichak KA, Kirkley KA (2017) Inflammatory Activation of Microglia and Astrocytes in Manganese Neurotoxicity. Adv Neurobiol 18:159–181.

Troche C, Aydemir TB, Cousins RJ (2016) Zinc transporter Slc39a14 regulates inflammatory signaling associated with hypertrophic adiposity. Am J Physiol Endocrinol Metab 310:E258–268.

Troyano-Rodriguez E, Mann S, Ullah R, Ahmad M (2019) PRRT1 regulates basal and plasticity-induced AMPA receptor trafficking. Mol Cell Neurosci 98:155–163.

Truett GE, Heeger P, Mynatt RL, Truett AA, Walker JA, Warman ML (2000) Preparation of PCR-quality mouse genomic DNA with hot sodium hydroxide and tris (HotSHOT). Biotechniques 29:52, 54.

Tuschl K, Clayton PT, Gospe SM, Mills PB (1993) Dystonia/Parkinsonism, Hypermanganesemia, Polycythemia, and Chronic Liver Disease.

Tuschl K, Mills PB, Parsons H, Malone M, Fowler D, Bitner-Glindzicz M, Clayton PT (2008) Hepatic cirrhosis, dystonia, polycythaemia and hypermanganesaemia--a new metabolic disorder. J Inherit Metab Dis 31:151–163.

Tuschl K, Clayton PT, Gospe SM, Jr., Gulab S, Ibrahim S, Singhi P, Aulakh R, Ribeiro RT, Barsottini OG, Zaki MS, Del Rosario ML, Dyack S, Price V, Rideout A, Gordon K, Wevers RA, Chong WK, Mills PB (2012) Syndrome of hepatic cirrhosis, dystonia, polycythemia, and hypermanganesemia caused by mutations in SLC30A10, a manganese transporter in man. Am J Hum Genet 90:457–466.

Tuschl K et al. (2016) Mutations in SLC39A14 disrupt manganese homeostasis and cause childhood-onset parkinsonism-dystonia. Nat Commun 7:11601.

Ubach J, Zhang X, Shao X, Sudhof TC, Rizo J (1998) Ca2+ binding to synaptotagmin: how many Ca2+ ions bind to the tip of a C2-domain? EMBO J 17:3921–3930.

Walden H, Muqit MM (2017) Ubiquitin and Parkinson’s disease through the looking glass of genetics. Biochem J 474:1439–1451.

Wang C, Ma Z, Yan DY, Liu C, Deng Y, Liu W, Xu ZF, Xu B (2018) Alpha-Synuclein and Calpains Disrupt SNARE-Mediated Synaptic Vesicle Fusion During Manganese Exposure in SH-SY5Y Cells. Cells 7.

Wang D, Zhang J, Jiang W, Cao Z, Zhao F, Cai T, Aschner M, Luo W (2017) The role of NLRP3-CASP1 in inflammasome-mediated neuroinflammation and autophagy dysfunction in manganese-induced, hippocampal-dependent impairment of learning and memory ability. Autophagy 13:914–927.

Wang WA, Groenendyk J, Michalak M (2012) Calreticulin signaling in health and disease. Int J Biochem Cell Biol 44:842–846.

Xu B, Shan M, Wang F, Deng Y, Liu W, Feng S, Yang TY, Xu ZF (2013) Endoplasmic reticulum stress signaling involvement in manganese-induced nerve cell damage in organotypic brain slice cultures. Toxicol Lett 222:239–246.

Yates AD et al. (2020) Ensembl 2020. Nucleic Acids Res 48:D682–D688.

Yin GN, Lee HW, Cho JY, Suk K (2009) Neuronal pentraxin receptor in cerebrospinal fluid as a potential biomarker for neurodegenerative diseases. Brain Res 1265:158–170.

Zeglam A, Abugrara A, Kabuka M (2018) Autosomal-recessive iron deficiency anemia, dystonia and hypermanganesemia caused by new variant mutation of the manganese transporter gene SLC39A14. Acta Neurol Belg.

Zhang J, Wang X, Vikash V, Ye Q, Wu D, Liu Y, Dong W (2016) ROS and ROS-Mediated Cellular Signaling. Oxid Med Cell Longev 2016:4350965.

Zhang Z, Yan J, Bowman AB, Bryan MR, Singh R, Aschner M (2019) Dysregulation of TFEB contributes to manganese-induced autophagic failure and mitochondrial dysfunction in astrocytes. Autophagy:1–18.

Zhi CN, Lai LL, Dou CS, Wang XH, Zhao P, Fu JL, Yao BY (2019) [The role of lysosomes in manganese-induced toxicity in SK-N-SH cells]. Zhonghua Lao Dong Wei Sheng Zhi Ye Bing Za Zhi 37:332–336.

Zhu G, Liu Y, Zhi Y, Jin Y, Li J, Shi W, Liu Y, Han Y, Yu S, Jiang J, Zhao X (2019) PKA- and Ca(2+)-dependent p38 MAPK/CREB activation protects against manganese-mediated neuronal apoptosis. Toxicol Lett 309:10–19.

Zofkova I, Davis M, Blahos J (2017) Trace elements have beneficial, as well as detrimental effects on bone homeostasis. Physiol Res 66:391–402.

Zou Y, Qing L, Zeng X, Shen Y, Zhong Y, Liu J, Li Q, Chen K, Lv Y, Huang D, Liang G, Zhang W, Chen L, Yang Y, Yang X (2014) Cognitive function and plasma BDNF levels among manganese-exposed smelters. Occup Environ Med 71:189–194.

