## Supplemental Material for "Loss of *slc39a14* causes simultaneous manganese deficiency and hypersensitivity in zebrafish"

### Supplementary Material

#### Supplementary Tables:

**Supplementary Table 1** List of differentially expressed genes identified by DeTCT and grouped by Mn toxicity (differentially expressed in $MnCl_2$ exposed siblings compared with unexposed siblings), Increased sensitivity (differentially expressed in $MnCl_2$ exposed mutants compared with unexposed siblings, but not differentially expressed in unexposed mutants compared to unexposed siblings or exposed siblings compared with unexposed siblings) and Genotype alone (differentially expressed in unexposed mutants compared with unexposed siblings). Genes highlighted in mustard are further discussed in the manuscript.

**Supplementary Table 2** List of enriched Gene Ontology (GO) terms.

**Supplementary Table 3** Statistical analysis of qRT-PCR data.

**Supplementary Table 4** List of enriched transcription factor motifs.

#### Supplementary Figures:

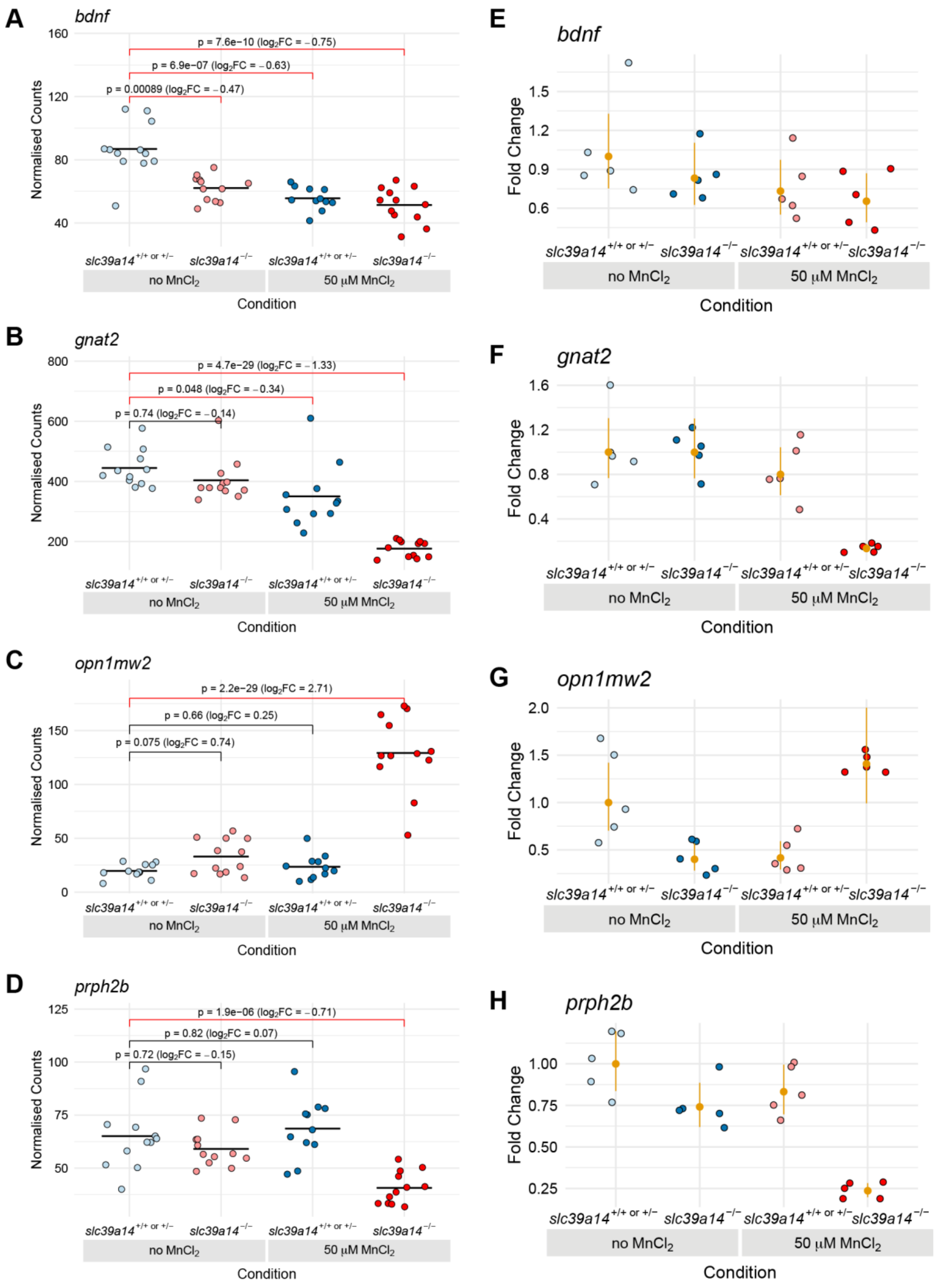


**Fig. S1. qRT-PCR produces consistent results with transcriptome sequencing.**
(A–D) Plots of the normalised counts for each sample for the genes *bdnf*, *gnat2*, *opn1mw2* and *prph2b*. Unexposed sibling embryos are light blue and $MnCl_2$ exposed ones are dark blue. Unexposed mutants are coloured light red and exposed mutants are dark red.
(E–H) Plots showing the qRT-PCR data for genes *bdnf*, *gnat2*, *opn1mw2* and *prph2b*. Values for individual samples are displayed as fold change relative to the mean value for unexposed siblings with the same colour scheme as in A–D. The mean and 95% confidence intervals for each condition are in orange.
